# Characterising Aromatic Side Chains in Proteins through the Synergistic Development of NMR Experiments and Deep Neural Networks

**DOI:** 10.1101/2024.04.01.587635

**Authors:** Vaibhav Kumar Shukla, Gogulan Karunanithy, Pramodh Vallurupalli, D Flemming Hansen

## Abstract

Nuclear magnetic resonance (NMR) spectroscopy has become an important technique in structural biology for characterising the structure, dynamics and interactions of macromolecules. While a plethora of NMR methods are now available to inform on backbone and methyl-bearing side-chains of proteins, a characterisation of aromatic side chains is more challenging and often requires specific labelling or ^13^C-detection. Here we present a deep neural network (DNN) named FID-Net-2, which transforms NMR spectra recorded on simple uniformly ^13^C labelled samples to yield high-quality ^1^H-^13^C correlation spectra of the aromatic side chains. Key to the success of the DNN is the design of a complementary set of NMR experiments that produce spectra with unique features to aid the DNN produce high-resolution aromatic ^1^H-^13^C correlation spectra with accurate intensities. The reconstructed spectra can be used for quantitative purposes as FID-Net-2 predicts uncertainties in the resulting spectra. We have validated the new methodology experimentally on protein samples ranging from 7 to 40 kDa in size. We demonstrate that the method can accurately reconstruct high resolution two-dimensional aromatic ^1^H-^13^C correlation maps, high resolution three-dimensional aromatic-methyl NOESY spectra to facilitate aromatic ^1^H-^13^C assignments, and that the intensities of peaks from the reconstructed aromatic ^1^H-^13^C correlation maps can be used to quantitatively characterise the kinetics of protein folding. More generally, we believe that this strategy of devising new NMR experiments specifically for analysis using customised DNNs represents a substantial advance that will have a major impact on the study of molecules using NMR in the years to come.

## Introduction

Nuclear Magnetic Resonance (NMR) spectroscopy is a ubiquitous technique in material science, chemistry, structural biology and clinical diagnosis. In bioscience, NMR provides unprecedented insight into functional motions (*1–7*) and non-covalent interactions (*8–10*) with atomic resolution. The technique therefore excellently complements AI-generated protein structures, *e.g.* from AlphaFold2, as well as structures obtained by cryo-electron microscopy (CryoEM) (*11–13*).

Over many decades, a series of developments that include advances in hardware, sample preparation, and novel NMR pulse sequences have steadily raised the ‘size-limits’ of proteins that can be studied using solution-state NMR. Specific advances include the introduction of per-deuteration (*14*), ^15^N-^1^H TROSY (*15*), and methyl-TROSY methods (*16*). Using these techniques, it is now possible to record amide ^15^N-^1^H and methyl ^13^C-^1^H correlation maps in megadalton sized proteins. However, studying functional side chains, such as charged or aromatic side chains, which are often present in enzymatic active sites and within interaction hotspots, are much more challenging.

We showed recently that employing ^13^C-detection allows for a characterisation of charged side chains, such as arginine and lysine, in proteins up to ∼40 kDa (*17*, *18*). For small proteins ^1^H-detected NMR methods are available to probe lysine and negatively charged side chains, which have provided insight into molecular recognition, salt-bridge, and hydrogen-bond formations (*19*, *20*). These experiments are often performed on uniformly ^13^C labelled proteins samples using constant-time (CT) experiments that eliminate the peak splitting arising due to homonuclear ^1^*J*_CC_ couplings in the indirect ^13^C dimension (*21*, *22*) to record high resolution [^13^C-^1^H] correlation maps at different backbone and side-chain sites.

Characterisation of aromatic side chains, on the other hand, has generally required specific labelling (*23–26*) because of non-uniform ^1^*J*_CC_ couplings and attenuation of signal due to substantial transverse relaxation during the constant-time period. There is therefore a clear need for improved methods to facilitate more detailed analysis of aromatic residues and their dynamics within proteins over a range of sizes of proteins to promote a greater understanding of how proteins function and interact.

Deep learning methods have had a substantial impact on all areas of science in recent years (*27*), solving key problems in biophysics and computational biology (*13*, *28*). Previous work from us and others have demonstrated applications of deep neural networks (DNNs) for transforming and analysing magnetic resonance data including analysing EPR DEER data (*29*), reconstructing non-uniformly sampled spectra, peak picking, and virtual homonuclear decoupling (*30–34*). Key to the success of these networks has been the ability to simulate an arbitrary amount of realistic training data (*29*, *34*), overcoming problems of overfitting and data bottlenecks that often beset these models. A shortcoming that exists in many existing DNNs in the field, however, is their inability to report reliable and quantitative uncertainties associated with the transformations.

In this work, we present a new DNN architecture, FID-Net-2, which uses data from a specially designed set of NMR experiments to not only reconstruct high resolution ^1^H-^13^C correlation maps of the aromatic side-chains in proteins, but also provide the uncertainty associated with the resulting spectra. The correlation maps generated by the DNN are free of the multiplet splittings and line broadenings that traditionally have degraded the quality of such spectra. We have validated the new DNN based methodology experimentally by accurately reconstructing high-resolution aromatic ^1^H-^13^C correlation spectra of the ∼20 kDa L99A mutant of T4 lysozyme (L99A-T4L) as well as the 40 kDa Maltose Binding Protein (MBP). Further, the utility of the new methodology is demonstrated by i) reconstructing high-resolution three-dimensional aromatic-methyl NOESY spectra to obtain aromatic ^1^H-^13^C assignments and ii) quantitating the peak intensities in the reconstructed high-resolution aromatic ^1^H-^13^C correlation maps recorded with varying exchange times to obtain the forward and reverse rate constants for the folding of the A39 FF domain from human HYPA/FBP11.

## Results

Due to variable ^1^*J*_CC_ couplings (*∼*55 to *∼*72 Hz) and fast ^13^C transverse relaxation, constant-time experiments are not routinely used to record high resolution ^13^C-^1^H correlation maps at various aromatic sites in proteins. Hence, we decided to develop a DNN to transform regular HSQC-like spectra, which contain multiplet splittings in the indirect (^13^C) dimension, into a high-resolution ^13^C-^1^H correlation map with sharp singlet peaks in the ^13^C dimension.

### Designing a pulse-sequence to aid recognition of the aromatic multiplet structure in proteins by the DNN

We have previously successfully trained the FID-Net (*31*) architecture to virtually decouple and enhance the resolution of ^13^C-^1^H correlation spectra reporting on the methyl-groups of large proteins (*35*). An initial attempt to use the same strategy for the aromatic region of ^13^C-^1^H correlation spectra of medium-to-large proteins was not satisfactory in our hands. We believe the reason for this is that the aromatic region of ^13^C-^1^H correlation spectra contains cross-peaks with different multiplet structures in the ^13^C dimension, whereas the methyl region essentially only contains doublets with a near uniform splitting of about ∼35 Hz. In the aromatic region, singlets are observed for histidine ^13^C^ε1^, doublets for tryptophan ^13^C^δ1^, and triplets for tyrosine and phenylalanine ^13^C^δ^ and ^13^C^ε^, respectively. Hence the DNN (or a human) cannot differentiate between two singlets with the same ^1^H chemical shifts separated by *∼*55 to *∼*72 Hz from a doublet, making it nearly impossible to train the DNN to perform a robust transformation between coupled and uncoupled spectra. Similarly, two doublets with the same ^1^H chemical shifts, ^1^*J*_CC_ couplings and chemical shifts differing by ^1^*J*_CC_ can be mistaken for a triplet. To facilitate a robust transformation by the DNN for resolution enhancement, we decided to take several steps. The first step was to design an NMR experiment that provides unique information about the multiplet structure of the cross-peaks so that the trained DNN can uniquely distinguish the multiplet structure of the cross-peak that it is transforming into a singlet. The DNN should then be able to avoid converting a doublet into two singlets or a triplet into two singlets.

The multiplet structure of the cross-peaks can be discerned by comparing two spectra: one corresponding to a normal ^13^C-^1^H HSQC spectrum and a second one in which the ^13^C-^13^C couplings have evolved for a small amount of time, τ_coup_ = 2.3 ms (∼1/6 ^1^*J*_CC_) (Figure 1a), in the indirect (^13^C) dimension. During the τ_coup_ delay, magnetisation arising from a singlet will not evolve while the two lines of the doublet will evolve with frequencies corresponding to ±*J*_CC_/2 and the time evolution of the two lines can be succinctly represented as {exp(− *i* π *J*_CC_ τ_coup_), exp(*i* π *J*_CC_ τ_coup_)}. Along similar lines, the two outer lines of a triplet will evolve with frequencies corresponding to ±*J*_CC_ while the inner line, that is of twice the hight, will not evolve its phase, and represented as {exp(− 2*i* π *J*_CC_ τ_coup_), 2, exp(2*i* π *J*_CC_ τ_coup_)}. Ignoring the effects of relaxation, spectra recorded with τ_coup_ = 0 and 2.3 ms will be indistinguishable from one another for a singlet. On the other hand spectra recorded with τ_coup_ = 2.3 ms from doublet and triplet sites will contain a combination of absorptive and dispersive lineshapes, while the τ_coup_ = 0 ms spectra only contains absorptive lineshapes. Ideal spectra calculated for the pair of experiments (red τ_coup_ = 0 ms (red); τ_coup_ = 2.3 ms (green)) are shown in Figure 1b for a singlet, in Figure 1c for a doublet and in Figure 1d for a triplet. Figure 1e shows a one-dimensional ^13^C slice extracted from ^1^H-^13^C datasets recorded on L99A-T4L using the complementary pair of experiments described. The slice originates from the ^13^C^δ2^ site of H31, where the spectrum recorded with τ_coup_ = 0.0 ms is in red and the one recorded with τ_coup_ = 2.3 ms is shown in green. The multiplet pattern arising from the regular spectra (red) in Figure 1e can arise either from two singlets or a doublet, but the spectrum recorded with τ_coup_ = 2.3 ms (green) that contains a combination of absorptive and dispersive lineshapes shows that it does not originate from two singlets (Figure 1b *vs.* 1e) but from a doublet (Figure 1c *vs* 1e). Along similar lines overlapping doublets can be distinguished from a triplet because the two components of the doublet evolve with frequencies of ±*J*_CC_/2 during the τ_coup_ = 2.3 ms delay while the components of the triplet evolve with a different set frequencies namely 0, ±^1^*J*_CC_ once again leading to different lineshapes in the spectra recorded with τ_coup_ = 2.3 ms. To summarise, the unique features, or pattern, generated by recording the second spectrum that incorporates evolution due to the ^13^C-^13^C coupling allows the DNN to identify the correct spin-system.

**Figure 1.**
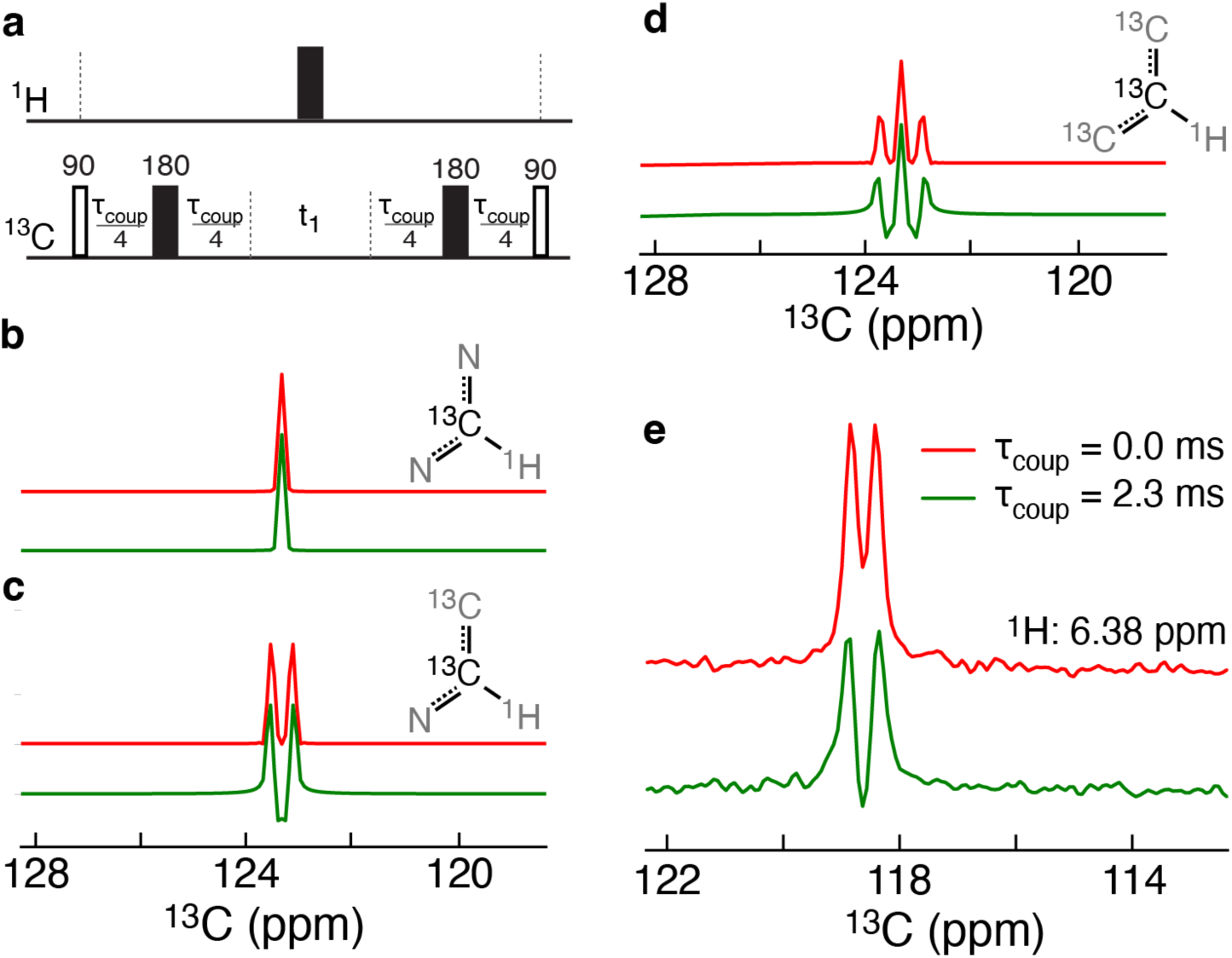
Encoding of unique features in ^13^C NMR spectra. (**a**) The core element of the pulse sequence that allows for evolution of the scalar couplings and thus encodes unique features of the multiplet structure. The chemical shift evolution time in the ^13^C dimension is denoted *t*_1_. Simulated 1D spectra showing the expected signals for a singlet (**b**), doublet (**c**) and a triplet (**d**) when the scalar couplings have been evolved for 0 ms (red) or 2.3 ms (green). *^1^J_CC_* was set to 70 Hz while the transverse relaxation rate was set 5 s^-1^. (**e**) One-dimensional ^13^C slices of a ^13^C,^1^H correlation spectrum on L99A-T4L recorded at a temperature of 298K and at a static magnetic field of 16.4 T. The slices are shown for the cross-peak arising from H31 ^13^C^δ2^-^1^H^δ2^ for τ_coup_ of 0.0 ms (red) and 2.3 ms (green).

### Training and assessing the performance of the FID-Net-2 DNN

To improve the spectral reconstruction from the two complementary datasets described above we made several key changes to the FID-Net architecture that we have devised previously. We name this new general architecture FID-Net-2. The main difference between the original FID-Net and the new FID-Net-2 architecture is that two complete 2D planes are processed within the architecture, as opposed to a sliding window of 1D spectra (Figure S1). Furthermore, FID-Net-2 outputs two sets of tensors (spectra), one output corresponding to the desired virtually decoupled and resolution-enhanced ^1^H,^13^C correlation spectrum, I(μ_H_,μ_C_), and a second tensor describing the uncertainty of the intensity for each point in the enhanced spectrum, α(μ_H_,μ_C_). The architecture is described in detail in Figure S1. Training a DNN such as FID-Net-2 requires a large amount of training data. For FID-Net-2 the training data consists of the complementary HSQC datasets with (2.3 ms) and without evolution due to ^1^*J*_CC_ couplings and a target high-resolution HSQC spectrum free of splittings in the ^13^C dimension. FID-Net-2 is then trained so that it learns to virtually decouple the desired high-resolution ^13^C-^1^H correlation map from the complementary HSQC datasets. The desired target high resolution HSQC spectrum free of splittings in the ^13^C dimension cannot be experimentally obtained from a uniformly ^13^C enriched sample and moreover would be infeasible to obtain for all the proteins required for training even if experimentally accessible. However, as we have now shown in multiple publications, it is now established that DNNs for transforming experimental NMR spectra can be trained on synthetic data. The FID-Net-2 model was trained on approximately 30×10^6^ sets of synthetically generated spectra.

The loss function (*Loss*_total_) developed for training FID-Net-2 includes three parts, *Loss*_total_ = *Loss*_1_ + *Loss*_2_ + *Loss*_3_. *Loss*_1_ corresponds to the traditional mean-square-error (MSE) between the target and predicted intensities. *Loss*_2_ was designed to ensure a gaussian distribution of the predicted uncertainties and *Loss*_3_ was designed to ensure that the uncertainties predicted agree with the RMSD between the target and predicted spectra. See materials and methods for a detailed description of the training procedure. Finally, it should be noted that FID-Net-2 can reconstruct high-resolution ^1^H-^13^C correlation maps from complementary HMQC or HSQC datasets because the same ^13^C chemical shift and the ^1^*J*_CC_ terms of the Hamiltonian are active during the *t*_1_ evolution period (^13^C dimension) in both of these experiments.

We initially assessed the performance of the trained FID-Net-2 model on sets of synthetic data, where the advantage is that the ground-truth is known. A summary of this assessment is shown in Figure 2. Figure 2a shows a representative example where FID-Net-2 is applied to a spectrum expected from an approximately 20 kDa protein at 298K. For such a case we expect about 50 cross-peaks and transverse relaxation rates of about 45 ± 20 s^-1^ in both the ^13^C and ^1^H dimensions. In contrast to other DNN transformations of NMR data, FID-Net-2 transforms the input and produces two outputs, that is, the desired correlation spectrum (middle) and the uncertainty associated with the transformation (right). Note that the input consists of two 2D planes, whereas only one is shown in Figure 2a.

**Figure 2.**
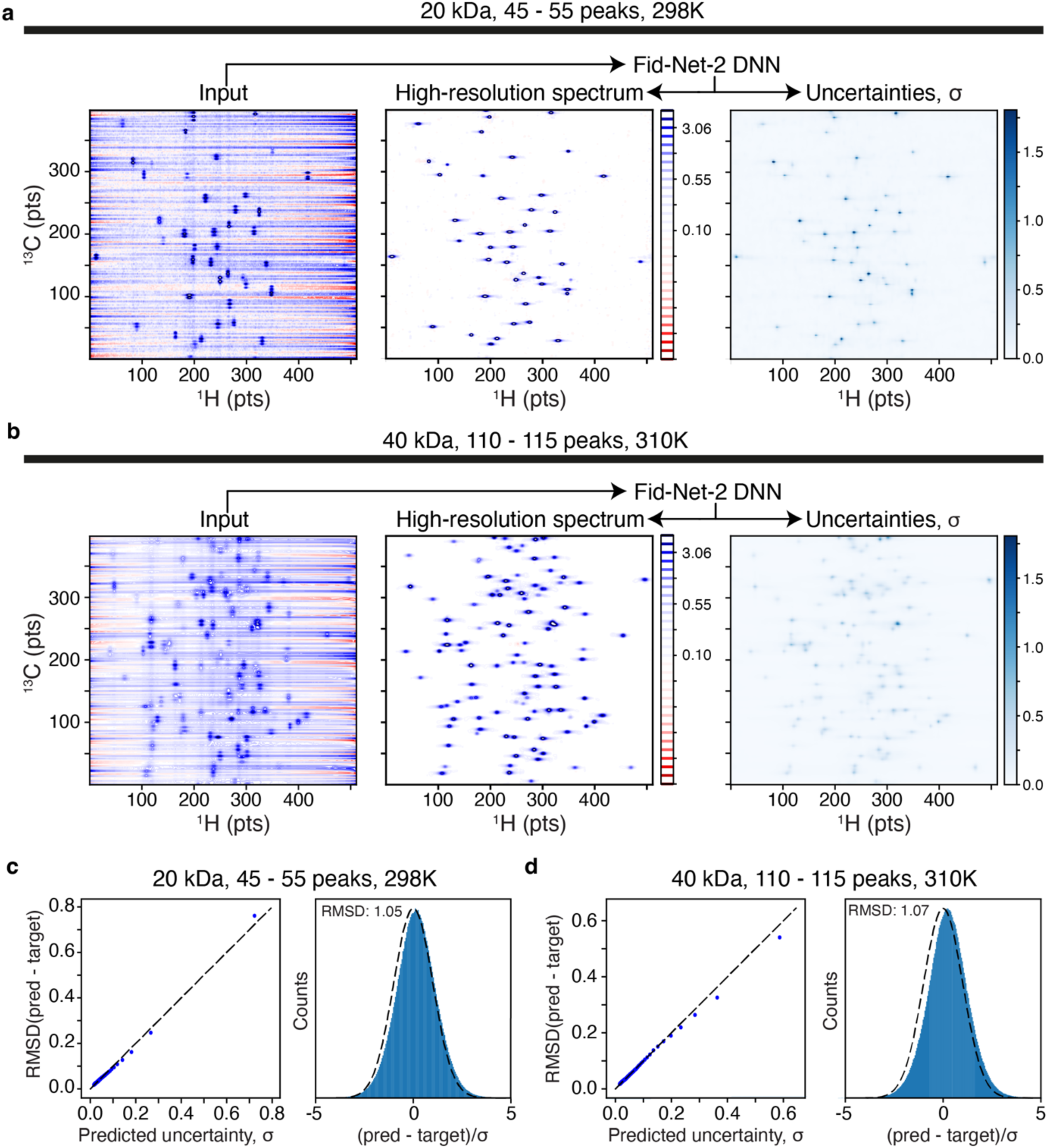
(**a**) Transformation with FID-Net-2 of randomly generated synthetic data corresponding to a 20 kDa protein (298K; 700 MHz). The transverse relaxation rates in the ^13^C and ^1^H dimensions were chosen from a random distribution with mean of 45 s^-1^ and standard deviation of 20 s^-1^. Other parameters match those in Table S1. (**b**) Transformation with FID-Net-2 of randomly generated synthetic data corresponding to a 40 kDa protein (310K; 700 MHz). The transverse relaxation rates in the ^13^C and ^1^H dimensions were chosen from a random distribution with mean of 95 s^-1^ and standard deviation of 20 s^-1^. Other parameters match those in Table S1. (**c,d**) Assessment of the predicted error, where to the left is the *χ*_i_ *v.s.* RSMD and to the right is a histogram of the calculated *χ*_i_ = (*pred*_i_ - *target*_i_)/σ_i_, showed a normal distribution with mean of nearly 0 and standard deviation of nearly 1. The plots in (**c)** and **(d**) are calculated over 10 random spectra, each with a *Loss*_1_ between 6.0×10^-3^ and 7.0×10^-3^, meaning these data are representing data amongst the worst 40%.

Firstly, it is seen that FID-Net-2 is able to eliminate the strong solvent signal to produce well-resolved spectra consisting of singlet cross-peaks. Of note is that the trained FID-Net-2 indeed produces point-by-point uncertainties, σ_i_, that match what is expected, as judged from a gaussian distribution of *Χ_i_* = (*target_i_* − *predicted_i_*)/*σ_i_*, and predicted σ_i_ that match the RMSD obtained from differences between predicted and target spectra (Figure 2c). Figure 2b shows an application of FID-Net-2 to a simulated spectrum of a larger protein with a molecular mass of about 40 kDa. For such a protein one expects about 110 cross-peaks in the aromatic region and transverse relaxation rates of about 95 ± 20 s^-1^. Again, the transformation of the input produces a clean well-resolved spectrum with predicted uncertainties that follow the desired criteria (Figure 2d). Effectively, Figures 2c,d shows that the implementation of *Loss*_2_ and *Loss*_3_ was successful.

One could argue that real experimental spectra potentially contain features, or artefacts, that have not been included in the training data, or that there is the potential that a future user will obtain data that contains artefacts that have not been included in the simulation data. Thus, we have not aimed to include every possible artefact that a future user might encounter in the training data, but instead show that the trained FID-Net-2 model is robust when transforming data that contains artefact not included in the training set. To test the robustness of FID-Net-2, and in particular its ability to produce reliable error estimates, we produced synthetic data where the common artefact of *t*_1_-noise encountered in NMR spectroscopy was included (Figure S2). Although *t*_1_-noise was not included in the training data in anyway FID-Net-2 reconstructed the desired spectrum from the input data and more importantly predicted uncertainties that are only slightly underestimated from the expected ones (Figure S2). Thus, although this is not a comprehensive analysis of all possible artefacts, one can expect that, when situations that have not been included during training are encountered, FID-Net-2 will report larger errors that agree with the uncertainty of the predicted spectrum.

### FID-Net-2 reconstructs high-resolution aromatic ^13^C-^1^H correlation maps from experimental data

Evaluations and assessments on synthetic data as shown above are important to judge the limitations of the trained FID-Net-2 model. However, it is by applying FID-Net-2 to real experimental data that we will truly understand its capabilities. Initially we recorded aromatic two-dimensional ^13^C-^1^H HSQC correlation spectra of the 18 kDa, L99A mutant of lysozyme from the phage T4 (*36*) (L99A-T4L) at 16.4 T (700 MHz), Figure S3. Apart from being relatively large compared to other proteins whose aromatic residues have been examined using NMR, L99A-T4L also exhibits conformational exchange that results in differential line-broadening further testing the ability of FID-Net-2 to reconstruct high-resolution spectra from coupled spectra. As expected, using a traditional Fourier transform to process the ^13^C-^1^H HSQC data results in ^13^C-^1^H correlation maps with multiplets in the ^13^C dimension that show substantial overlap (Figure 3a). In contrast, when the complementary pair (with and without the coupling delays) of ^13^C-^1^H HSQC datasets are processed using the FID-Net-2 model, a well-resolved spectrum of high quality is obtained. Furthermore, the produced uncertainties are clearly not uniformly distributed over the spectrum as is the case for thermal noise processed with a linear Fourier transformation. It is well-known that DNNs produce mappings that are highly non-linear and one cannot therefore simply assess the performance, or accuracy, from the RMSD of a transformed spectrum in an area without cross-peaks, which is custom for standard processed spectra. The produced uncertainties in Figure 3c clearly show that the uncertainties are centred around strong cross-peaks and near highly overlapped peaks. The aromatic ^13^C-^1^H correlation maps reconstructed by FID-Net-2 from datasets with differing coupling delays are both better resolved and contain more signal compared to constant-time HSQC spectra (Figure S4).

**Figure 3.**
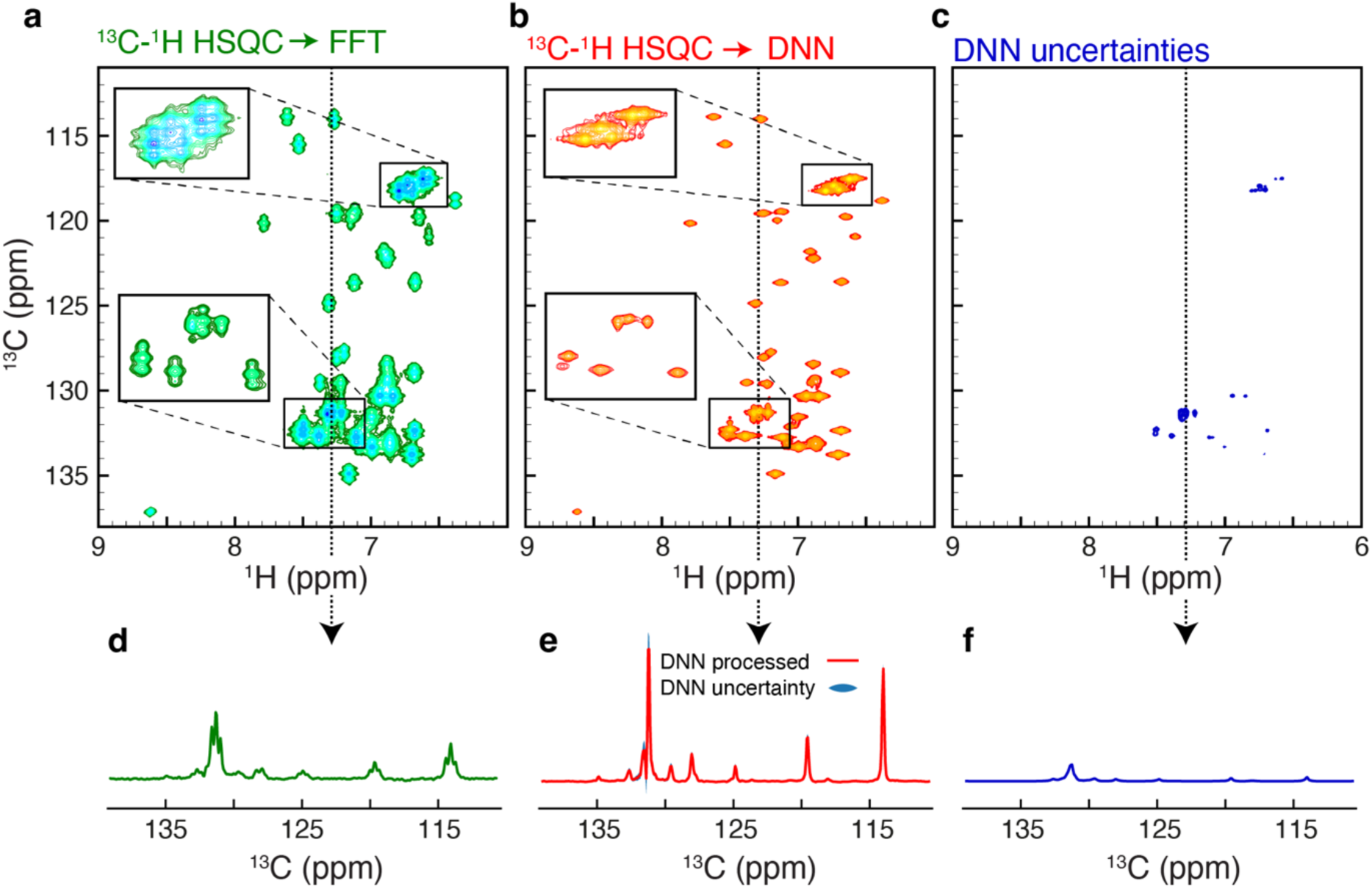
Transformation of experimental spectra of L99A-T4L. (**a**) ^13^C-^1^H HSQC spectrum reporting on the aromatic region of the 18 kDa L99A-T4L (298K; 700 MHz). Correlations with different coupling multiplicity are clearly visible which leads to severe overlap in this medium size protein. (**b**) The high resolution ^1^H-^13^C map reconstructed by FID-Net-2 from two ^13^C-^1^H HSQC spectra, recorded with τ_coup_ = 0.0 and 2.3 ms does not contain the multiplets seen in (**a**) leading to significantly lower overlap. (**c**) The uncertainty in the intensities of the reconstructed spectrum (**b**) predicted by FID-Net-2. (**d, e, f**) one-dimensional representative slices of the spectra in **a**, **b**, and **c**, respectively.

Having evaluated the trained FID-Net-2 model on synthetic data, including synthetic data with *t*_1_-noise, as well as on good-quality experimental data, we sought to further assess how the trained model behaves when the data contains artefacts that are not included in the training data. We did so experimentally by deliberately mis-setting the *Z*_1_ and *Z*_2_ shims of the NMR spectrometer to create an inhomogeneous field and thus create lineshapes that deviate dramatically from the Lorentzian lineshapes used for training (Figure S5). For L99A-T4L we recorded ^13^C-^1^H HSQC correlation spectra with optimal shimming and with non-optimal shimming and subsequently compared peak-intensities and peak-positions, in line with the NUScon criteria (*37*). Excellent correlations are obtained both for peak positions and intensities (Figure S5) showing that FID-Net-2 can robustly reconstruct spectra from experimental data recorded under suboptimal conditions.

### Applications to larger proteins: FID-Net-2 reconstructs the high-resolution aromatic ^13^C-^1^H correlation map of 40 kDa *E. coli* Maltose Binding Protein

Recording high resolution aromatic ^13^C-^1^H correlation maps for large proteins remains a challenge due to the short ^13^C transverse relaxation times that make constant-time HSQC spectra very insensitive. The HMQC spectrum recorded on 40 kDa *E*. *coli* Maltose Binding Protein in complex with β-Cyclodextrin (MBP) at 310K contains few resolved correlations (Figure 4a) and a large number of correlations are severely overlapped due to *^1^J*_CC_ splittings in the indirect dimension. The ^1^H-^13^C correlation map reconstructed by FID-Net-2 however is much better resolved, once again demonstrating the efficacy of FID-Net-2 at reconstructing high-resolution aromatic ^1^H-^13^C correlation maps. We have chosen to use HMQC rather than HSQC type datasets as they are about 10% more sensitive (see Figure S6). The NOESY based strategy described below can in principle be used for the assignment of the correlations in Figure 4b but this is beyond the scope of this work.

**Figure 4.**
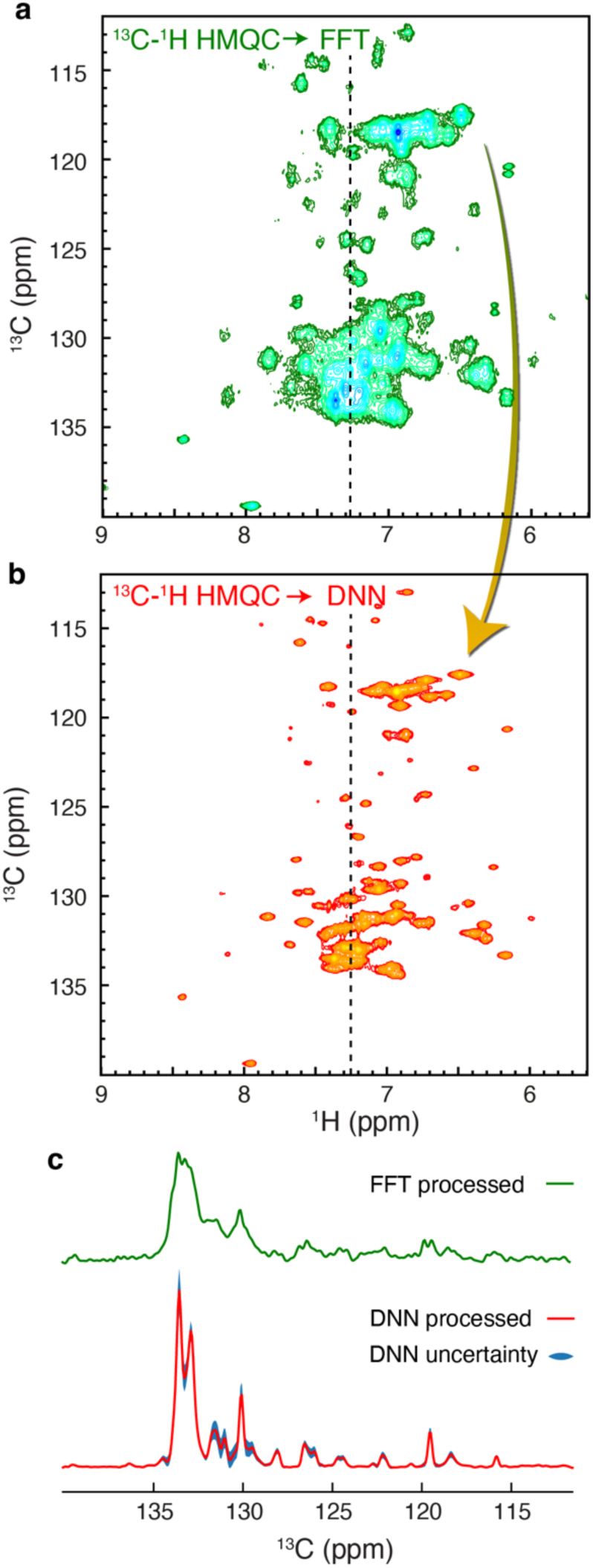
Transformation of experimental spectra of the 40 kDa MBP. (**a**) ^13^C-^1^H HMQC spectrum reporting on the aromatic region of the 40 kDa MBP, recorded at 310K and at 700 MHz. Substantial overlap is observed with few resolved cross-peaks. (**b**) Processing with the FID-Net-2 model of two ^13^C-^1^H HMQC spectra, recorded with τ_coup_ = 0.0 and 2.3 ms. Many well-defined cross-peaks are observed, and the overlap is substantially less than in **a**. (**c**) A one-dimensional slice of the input ^13^C-^1^H HMQC spectrum is compared with the corresponding one-dimensional slice of the output from FID-Net-2. The uncertainties predicted by the DNN model are shown as a blue filled area.

Using the 40 kDa MBP protein, with substantial peak overlap, we further assessed the FID-Net-2 mapping and the estimation of uncertainties. In summary, we recorded two sets of spectra, one with low signal-to-noise (8 scans) and one set with high signal-to-noise ratio (128 scans). Since these spectra were recorded on the same sample using the same NMR spectrometer (700 MHz; 310K), one expects that the signal intensities are proportional and that any deviations are captured by the uncertainties predicted by the trained FID-Net-2 model. Figure S7 shows an excellent correlation between the two transformed datasets, and it also shows that the deviations are well captured by the predicted uncertainties, thus providing further evidence that the trained FID-Net-2 model transforms the data accurately, even noisy data, and also produces quantitative uncertainties.

### Exploiting FID-Net-2 to obtain aromatic ^1^H-^13^C assignments from NOESY experiments

Obtaining aromatic ^1^H and ^13^C assignments in medium size proteins is challenging because HSQC-NOESY type spectra have poor resolution in the aromatic ^13^C dimension due to ^1^*J*_CC_ couplings, while the CT-HSQC-NOESY spectra suffer from poor signal-to-noise due to the short transverse relaxation times of aromatic ^13^C nuclei. FID-Net-2 provides a ready solution to the problem. In order to assign the chemical shifts of the aromatic ^13^C-^1^H spectrum of L99A-T4L, we recorded ^13^C_Methyl_-^13^C_Aromatic_-^1^H_Aromatic_ and ^1^H-^13^C_Aromatic_-^1^H_Aromatic_ three-dimensional NOESY spectra (Figure S8) and processed these with FID-Net-2 in the ^13^C_Aromatic_-^1^H_Aromatic_ dimensions. A summary of these spectra and the chemical shift assignment procedure that utilises ^13^C,^1^H methyl assignments are shown in Figure 5. Figure 5a highlights how the uncertainties in intensity provided by FID-Net-2 aid in analysing the NOESY spectra. Cross-peaks with uncertainties that are as large as the signal intensities should be very carefully assessed, whereas cross-peaks (even weak ones), with small uncertainties can be confidently interpreted. Based on a previous ^13^C,^1^H methyl assignment (*38*), these two spectra were sufficient to assign the correlations seen in the high-resolution aromatic ^13^C,^1^H correlation map (Figure S9).

**Figure 5.**
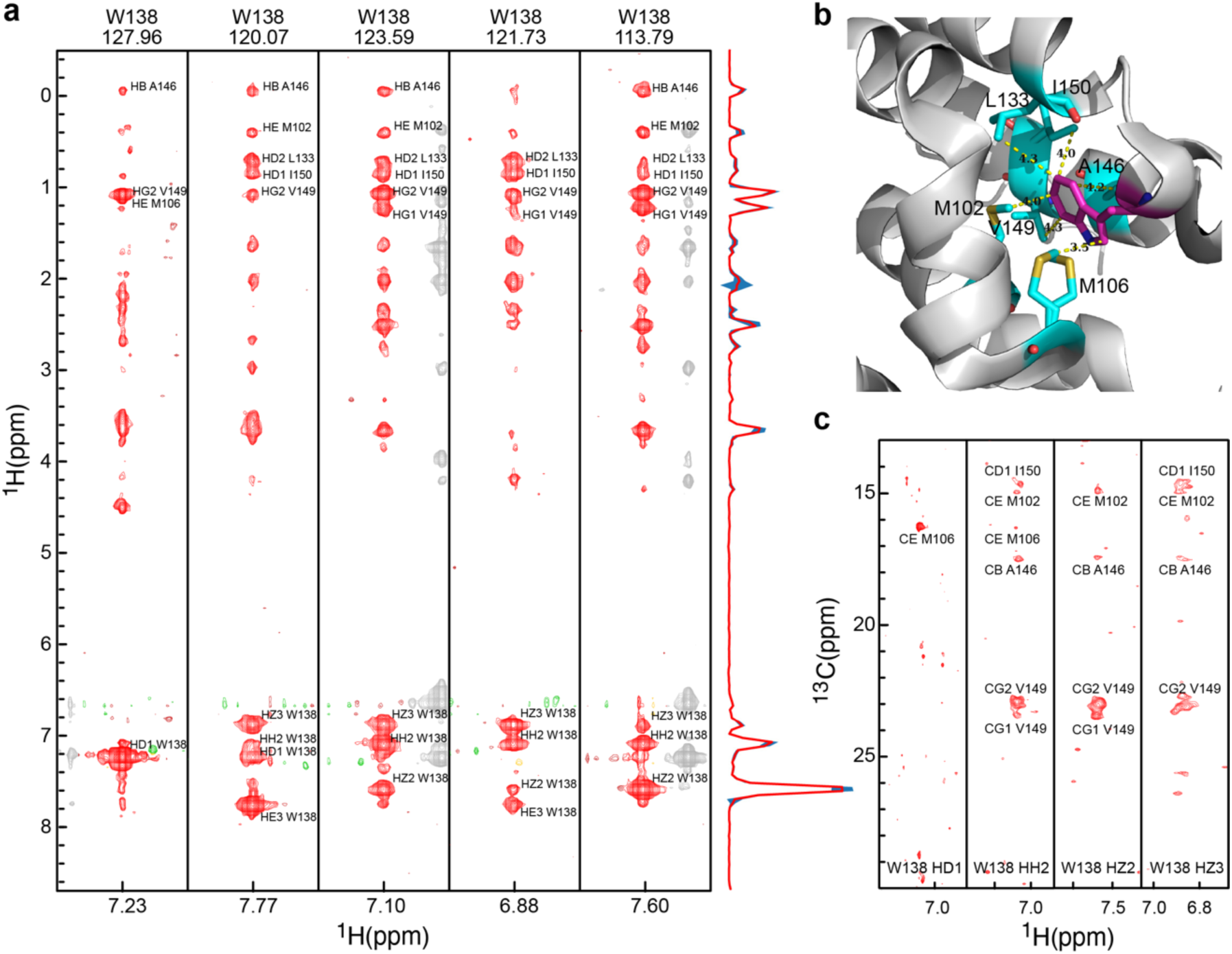
Aromatic ^1^H-^13^C assignments from NOESY spectra reconstructed using FID-Net-2. (**a**) Strips from the ^1^H-^1^H planes of the 3D ^1^H-^13^C_Aromatic_-^1^H_Aromatic_ NOESY spectrum of L99A-T4 Lysozyme (25 °C; 700 MHz) used for the assignment of Trp 138. **(b)** The residue Trp 138 is highlighted as magenta sticks on a cartoon representation of the T4 Lysozyme structure [PDB ID: 3dmv] (*39*). The residues in close proximity to Trp 138 are shown in cyan sticks and their distances from the aromatic side-chain of Trp 138 are also shown in the figure. **(c)** Strips from the ^1^H-^13^C planes of the 3D ^13^C_Methyl_-^13^C_Aromatic_-^1^H_Aromatic_ NOESY spectrum of L99A-T4 Lysozyme (25 °C; 700 MHz) focussing on Trp 138. The structure of the protein was used to identify aromatic and methyl protons that are close to one another, following which the complementary pair of 3D NOESY spectra that contain cross peaks between aromatic and methyl protons that are proximal to one another was used to assign the aromatic ^1^H and ^13^C resonances. FID-Net-2 was used to process the ^13^C_Aromatic_-^1^H_Aromatic_ dimensions.

### Quantitative characterisation of protein dynamics using FID-Net-2

Previous DNNs devised to transform NMR spectra were not quantitative with respect to the intensities of cross-peaks (*35*) and were not useful to study chemical exchange, characterise binding or other studies where accurate peak intensities are necessary. FID-Net-2 was however trained to be quantitative in this regard. To exploit this aspect of FID-Net-2, we recorded longitudinal exchange (*40*) spectra (EXSY/ZZ exchange) on the A39G mutants of the FF domain (A39G-FF), Figure 6. The aromatic ^13^C,^1^H chemical shift assignment of A39G-FF was obtained using the 3D NOESY spectra described above, Figure S10. A39G-FF exchanges slowly between the folded state and the unfolded state (*41*) and the addition of a small amount of urea (1 M) increases the unfolded state population giving rise to two sets of peaks in NMR spectra. As seen in Figure 6b, the FID-Net-2 transformed ^13^C,^1^H correlation map clearly shows the two sets of cross-peaks reporting on the exchange between the folded and unfolded states of A39G-FF. A least-squares analysis of the data provided the exchange rate (*k*_ex_) and the population of the unfolded species (*p*_U_). To assess the quality of the data, we also recorded ^15^N,^1^H ZZ exchange spectra and obtained an exchange rate and a population (*k*_ex_ = 4.1 ± 0.2 s^-1^ and *p*_U_ = 38.7 ± 0.8%.) in agreement with those obtained from the FID-Net-2 transformed spectra (*k*_ex_ = 3.4 ± 0.3 s^-1^ and *p*_U_ = 36.3 ± 1.7%) thus experimentally demonstrating that spectra transformed with FID-Net-2 can be used for quantitative analyses.

**Figure 6.**
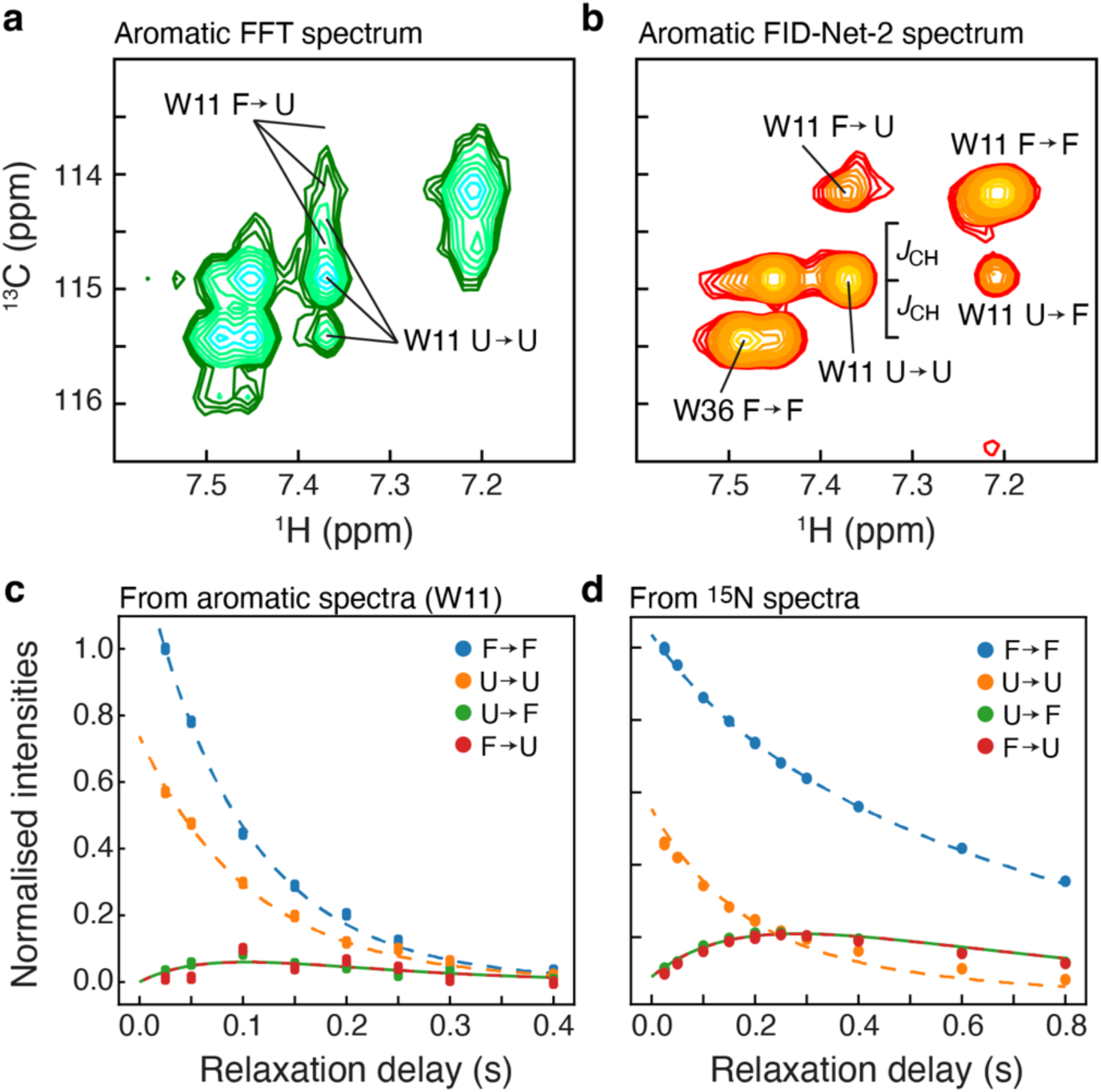
Transformations with FID-Net-2 are quantitative. Regular ^13^C-^1^H correlation map (**a**) and the FID-Net-2 reconstructed ^13^C-^1^H correlation map from a ZZ exchange (*T_EX_* = 150 ms) experiment reporting on the aromatic region of the 7 kDa the A39G mutant FF domain in the presence of 1M Urea (275K; 600 MHz). Both spectra contain peaks arising from the folded (F) as well as the unfolded (U) state of the protein. The regular (FFT) spectrum (**a**) is severely overlapped while the FID-Net-2 reconstructed spectrum (**b**) is much better resolved allowing one to identify both diagonal (F ➝ F and U ➝ U) as well as exchange cross-peaks (F ➝ U and U ➝ F) arising from the ^13^C^ζ2^-^1^H^ζ2^ site in W11. (**c**) Intensities extracted from (**b**) for various *T_EX_* delays were analysed using the standard Bloch-McConnnell formalism (*42*) to obtain the exchange parameters. The dashed lines are drawn using the best fit parameters (*k*_ex_ = 3.39 ± 0.32 s^-1^ and *p*_U_ = 36.3 ± 1.7 %). (**d**) Intensities extracted a ^15^N ZZ exchange experiment on the same sample, for diagonal (F ➝ F and U ➝ U) and well as exchange peaks (F ➝ U and U ➝ F). The dashed lines are drawn using the best fit exchange parameters (*k*_ex_ = 4.08 ± 0.17 s^-1^ and *p*_U_ = 38.7 ± 0.8 %).

## Discussion

Being able to characterise the regulation, interactions, and dynamics of medium and large proteins in solution is paramount to understanding molecular functions. To that end, it is imperative to have tools to characterise aromatic side chains in proteins that are critical reporters of function because these sites are often located in interaction hot spots, involved with substrate binding, regulation and catalysis.

Specific isotopic labelling (*23–25*, *43*, *44*), has been one of the only means to characterise aromatic residues in medium-sized proteins. However, these labelling schemes limit the number of probes available and require the use of specific precursors that often lead to reduced protein yield of the samples. Here we presented an attractive alternate method to characterise functional aromatic residues in medium-sized proteins, wherein a pair of complementary ^1^H-^13^C datasets recorded using a uniformly ^13^C-isotopically enriched protein sample are processed with the FID-Net-2 model to obtain the desired high-resolution aromatic ^13^C-^1^H correlation map. It is important to note that this methodology, based on processing with a deep neural network, offers simultaneous access to all the ^13^C-^1^H spin-pairs in all the aromatic side chains in the protein and does not require specifically labelled samples. The FID-Net-2 network architecture is itself providing a new way to transforming NMR spectra using DNNs, because it not only produces resolution enhanced spectra, but also provides a good estimate of the uncertainty in the intensities of these spectra. We have exploited these abilities of FID-Net-2 by obtaining chemical shift assignments (L99A-T4L) and characterising chemical exchange (A39G-FF). We believe that our new methodology will allow for a general and easy characterisation of functional aromatic side chains in medium-sized proteins.

Two major developments contribute to the success of FID-Net-2: i) the design of new NMR experiments with the sole goal of aiding the DNN and ii) training the DNN to estimate uncertainties of the transformed spectra. Datasets with τ_coup_ set to 2.3 ms are recorded solely to provide unique features for the DNN to analyse. Due to ^1^*J*_CC_ evolutions during τ_coup_, spectra obtained from such datasets will contain dispersive components in the ^13^C dimension making them unappealing to a human NMR spectroscopist, but nonetheless useful to the DNN that utilises the information present in such datasets to reconstruct high resolution ^1^H-^13^C correlation maps. The uncertainties estimated by FID-Net-2 are crucial to both applications presented here. Knowledge of the uncertainties was critical for both identifying ‘valid’ cross-peaks in the NOESY spectra for the purposes of assignment and for obtaining kinetic parameters from the variation of cross-peak intensities as a function of mixing time. As with other convolutional neural networks, it is likely that the trained FID-Net-2 model presented in this study can be re-trained to transform other types of spectra.

It is now clear that processing and transforming NMR spectra with DNNs is a powerful tool. However, we believe that to truly exploit the potential of DNNs in NMR, it is not enough to just devise new DNNs that transform existing experimental data, but to devise new experiments specifically for the DNNs to exploit as we have done here. Concomitantly developing DNNs and experimental methods will in the future to come allow for new insights, in AI-assisted NMR spectroscopy and likely also in other related scientific fields.

## Materials and Methods

### The FID-Net-2 architecture

Our aim was to develop a DNN to map ^13^C-^1^H correlation NMR spectra reporting on the aromatic region of uniformly ^13^C-labelled proteins into spectra of high resolution. Standard ^13^C-^1^H spectra of uniformly labelled proteins are affected by one-bond ^13^C-^13^C homonuclear scalar couplings, line broadenings, and residual solvent signals. The developed DNN will therefore need to (*i*) virtually decouple the multiplet structures arising from the homonuclear couplings, (*ii*) generally enhance the resolution, and (*iii*) remove solvent signals. Finally, (*iv*) we also require that the DNN is able to predict the accuracy with which it does the mapping, which means that the DNN provides point-by-point uncertainties α (μ_1H_,μ_13C_), of the predicted output I(μ_1H_,μ_13C_). As noted in the main text and Figure 1, two input spectra are required in order for this transformation to be robust. It should be noted that the mapping performed by the developed DNN will not increase the information in the provided data, but will combine the information in the two input spectra and generate a spectrum that is of high resolution and easily interpretable by the end-user spectroscopist.

To achieve the above requirements for the DNN, the previous FID-Net architecture (*31*) was substantially altered in several ways, including, (*i*) full 2D planes are transformed as opposed to using a sliding window, (*ii*) both the ^13^C and ^1^H dimensions are processed within the same architecture, (*iii*) a refinement step in the frequency domain was included in the end, and (*iv*) uncertainties are also predicted. Of note is that the last layer of FID-Net-2 produces a tensor of size (512,400,2), where the first (512,400) plane is the ^1^H-^13^C resolution enhanced spectrum and the second (512,400) plane is the confidences. A sigmoidal activation, 1/(exp(-x)+1), is used to ensure that the confidences take values between 0 and 1. Standard deviations are calculated from the confidence, conf, by:

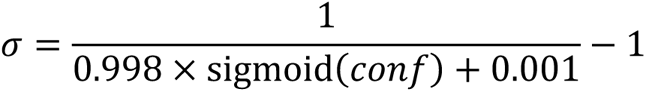

Finally, the predicted spectrum and the predicted uncertainties, *σ*, are convolved with a sine-bell window function, with offset of 0.4π, before calculating the losses. The architecture is detailed in Figure S1.

### Synthetic spectra for training FID-Net-2

The FID-Net-2 DNN was trained exclusively on synthetic data, summarised in Figure 2, and subsequently evaluated on synthetic data and experimentally acquired from protein samples. The resolution in the ^13^C dimension was enhanced both with virtual decoupling and by decreasing the effective transverse relaxation rate. When decreasing the effective transverse relaxation rate, care must be taken, so that the DNN does not generate artefacts from very broad features in the spectrum. We found that halving the effective relaxation rate worked well in the ^13^C-dimension, that is, *R*_2,tar_ = 0.5 *R*_2,inp_, where the input rates, *R*_2,inp_, were randomly generated from a normal distribution with mean of 50 s^-1^ and standard deviation of 20 s^-1^ and *R*_2,tar_ is the target transverse relaxation rate. The multiplet structures of the ^13^C-^13^C couplings in the input spectrum were simulated by generating two sets of coupling constants, *J*_1,C_ and *J*_2,C_, that were each drawn from a normal distribution with mean of 63 Hz and standard deviation of 10 Hz. Subsequently 20% of *J*_1,C_ and 20% of *J*_2,C_ were set to zero, which results in 64% triplet structures, 4% singlet structures, and 32% doublet structures. To simulate non-weak couplings, roofing effects were added by multiplying the FID in the ^13^C dimension by {cos(*πJ*_1,C_*t*_1_) + *ϱ*_1_ *i* sin (*πJ*_1,C_*t*_1_)} } {cos(*πJ*_2,C_*t*_1_)} + ϱ *i* sin(*πJ*_2,C_*t*_1_)}. Here *ρ*_1_ and *ρ*_2_ are factors to include roofing effects and these are both drawn from a normal distribution with mean of 0 and standard deviation of 0.08. Each spectrum contained between 40 and 200 cross-peaks, with chemical shifts uniformly distributed along the ^13^C-dimension. The ^1^H chemical shift were generated to increase the overlap of cross-peaks. Firstly, initial ^1^H chemical shifts ƍ_H_^0^ were drawn from a normal distribution with mean of 0 ppm and standard deviation of SW/4. Subsequently, to increase the overlap, the final ^1^H chemical shifts were calculated using the following empirical equation, which also ensures that cross-peaks are not on the edge of the spectrum in the ^1^H dimension:

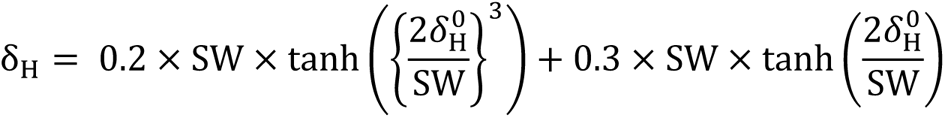

In the ^1^H dimension, the input simulated data included ^1^H-^1^H homonuclear couplings. Similar to the ^13^C dimension, two sets of coupling constants were generated *J*_1,H_ with an average of 8 Hz and a standard deviation of 2 Hz and *J*_2,H_ with an average of 4 Hz and a standard deviation of 2 Hz; 10% of *J*_1,H_ were set to zero and 50% of *J*_2,H_ were set to zero. Roofing effects, that is non-week couplings, were simulated in the same way as for the ^13^C-dimension. Solvent signals were simulated in the ^1^H frequency domain as

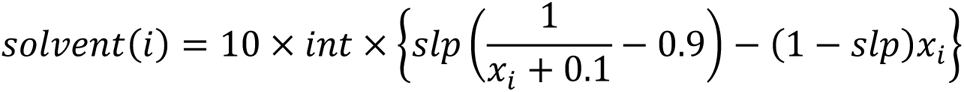

where *int* is a random number drawn from the same normal distribution used to assign peak intensities, *slp* is a random number between 0 and 1 (uniform). When the 1D ^1^H frequency-domain spectrum contains *N* points, then *x*_i_ is 0/*N*, …, (*N*-1)/*N*. Half of the solvent residuals were inverted in the ^1^H dimension, such that the DNN learned to deal with solvent signals from both the left and the right side of the spectrum. Finally, the residual solvent signal generated in frequency domain was Fourier transformed to generate the solvent signal in the time domain, which was added to the synthetically generated random spectrum.

The FID-Net-2 model was trained on a diverse range of NMR parameters (Table S1) and so can be used without need for further retraining and the approach can be used with standard ^1^H-^13^C HSQC or HMQC pulse sequences.

### Training the FID-Net-2 architecture with synthetic spectra

The FID-Net-2 model was trained on approximately 30×10^6^ sets of spectra, where one set consisted of a target 2D spectrum (*target*) and two input spectra, without coupling evolution, *input*_no-coup_, and with 2.3 ms coupling evolution in the ^13^C dimension, *input*_coup_. Briefly, chemical shifts were randomly distributed in the ^13^C dimension, while more condensed in the ^1^H dimension to mimic increased overlap. For the input spectra, we also added random gaussian noise and a solvent signal akin to a residual water signal. A maximum of 200 cross-peaks were generated. All training parameters are provided in Table S1. The DNN model, Figure S1, was developed and trained using the TENSORFLOW 2.11 library (*45*) with the KERAS (*46*). As mentioned in the text specialised loss functions were used to train the network with the total Loss (*Loss*_total_ = *Loss*_1_ + *Loss*_2_ + *Loss*_3_). *Loss*_1_ corresponds to the traditional idea of minimising the difference between the predicted output spectrum and the target spectrum, whereas *Loss*_2_ is restraining a Gaussian distribution of the predicted errors, and *Loss*_3_ is restraining the calculated uncertainties to match the RMSD between the predicted and target spectrum, over 200 linear bins. Specifically, the function for *Loss*_1_, also referred to as mean-square-error (MSE) is simply defined as:

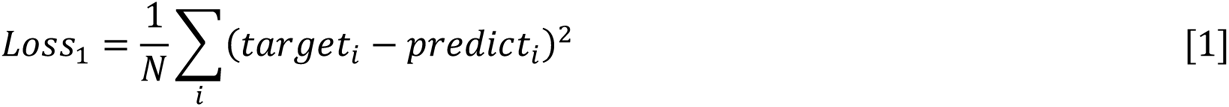

Where the sum is over all points in the spectrum and *N* is the total number of points in the 2D plane (400 × 512). The losses *Loss*_2_ and *Loss*_3_ were designed specifically for Fid-Net-2. For *Loss*_2_, a value *Χ_i_* was first calculated as

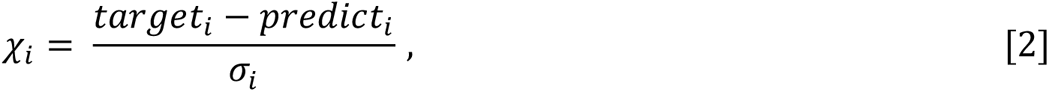

where *σ_i_* is the predicted error (output from FID-Net-2). Our goal was to have *Χ_i_* follow a standard gaussian distribution with zero mean and standard deviation of 1. To achieve this, the 1/2^th^, 1^st^, 2^nd^, 3^rd^, and 7/2^th^ momenta of *Χ_i_* were restrained as follows,

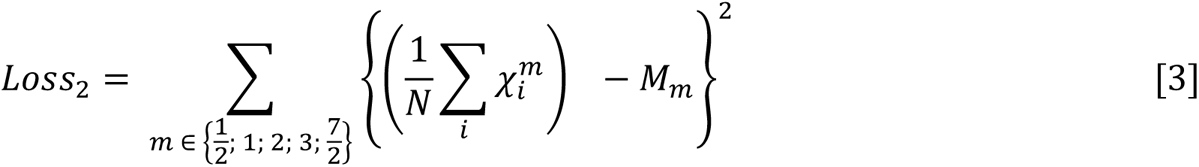

where 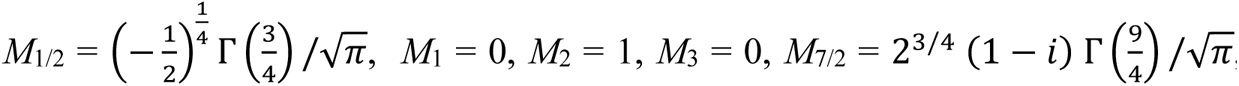, and Γ() is the gamma function.

For the calculation of *Loss*_3_, the predicted errors, *σ*_i_, were binned into 200 bins (linear), with the bins equally spaced between 0 and max(*σ*_i_). Within each of these 200 bins, the average of the *σ*_i_ was calculated and restrained to be equal to the RMSD between the predicted points and the target points, for points corresponding to this bin. Specifically,

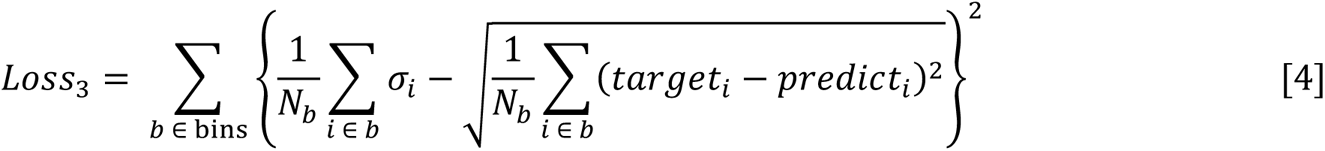

The model was trained to a total loss of 8.7×10^-3^ and *Loss*_1_ of *ca.* 5×10^-3^. Below, the trained model is first assessed on synthetic data and subsequently we evaluate the model on a series of experimental data.

The ADAM (*47*) optimiser was used for training with a learning rate that changed throughout the training, *β*_1_=0.9, *β*_2_=0.98, and ε=10^-9^. Mini-batching was used with 4 set of spectra in each mini-batch and the weights saved for every 2000 batches. The learning rate, *lr*, was calculated as

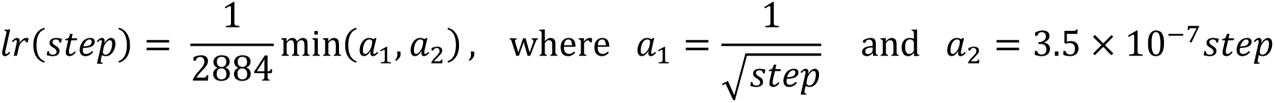

The parameter, *step*, is a counter for batches used in training. Thus, after an initial warm-up period, the largest learning rate used is about 2.45×10^-6^, whereafter the learning rate decays. The FID-Net-2 architecture was trained on the NMRBox facility (*48*) using two Nvidia A100 GPUs.

### Assessment using synthetic data

Once trained, the performance of the trained FID-Net-2 model was initially evaluated on synthetically generated data, as shown in Figure 2. Two independent assessments were made, one to represent a protein of about 20 kDa and one to represent a protein of about 40 kDa. Parameters used to generate the synthetic spectra were the same as those used for training, Table S1, except that for the 20 kDa protein only between 45 and 55 cross-peaks were generated with *R*_2_(^1^H) and *R*_2_(^13^C) both drawn from a normal distribution with mean of 45 s^-1^ and standard deviation of 20 s^-1^. For the 40 kDa protein 110-120 cross-peaks were generated with *R*_2_(^1^H) and *R*_2_(^13^C) both drawn from a normal distribution with mean of 95 s^-1^ and standard deviation of 20 s^-1^.

### NMR Samples

All three [U -^15^N^13^C] protein samples were prepared by over expressing the proteins in *E. coli* BL21(DE3) cells transformed with the appropriate plasmids and grown in M9 medium supplemented with 1 g/L of ^15^NH_4_Cl and 3 g/L of ^13^C-glucose as nitrogen and carbon sources respectively. L99A-T4L (*38*), A39G-FF (*41*) and MBP (*49*) were all purified as described previously.

The L99A-T4L sample consisted of ∼1 mM [U -^15^N^13^C] protein dissolved in 50 mM sodium phosphate, 25 mM NaCl, 2 mM EDTA, 2 mM NaN_3_, ∼99% D_2_O, pH 5.5 buffer. The A39G-FF samples consisted of ∼1 mM [U -^15^N^13^C] protein dissolved in 50 mM sodium acetate, 100 mM NaCl, 2 mM EDTA, 2% D_2_O pH 5.7 buffer. The MBP samples consisted of ∼0.5 mM [U -^15^N^13^C] protein dissolved in 20mM sodium phosphate, 1mM EDTA, 2mM β-cyclodextrin, 99% D_2_O, pH 6.5 buffer.

### NMR Experiments

#### Two-dimensional ^13^C-^1^H correlation spectra

The 2D HSQC and HMQC datasets of L99A-T4L (Figure 3) and MBP (Figure 4) used as input for FID-Net-2 were recorded on a 1 mM (L99A-T4L) or 0.5 mM (MBP) uniformly [^13^C,^15^N]-labelled samples using the pulse sequences described in Figure S3a,b, with τ_coup_ = 0.0 ms and 2.3 ms, on a Bruker 700 MHz Avance III spectrometer equipped with Z-gradient triple-resonance TCI cryoprobe. The data was acquired with 768 and 200 (L99A-T4L) or 128 (MBP) complex points in the ^1^H and ^13^C dimensions, respectively, with spectral widths of 10000 Hz and 5000 Hz. The datasets were recorded as pseudo-3D spectra. An interscan delay of 1 s was used.

The constant-time experiments (Figure S4) were recorded on a 1 mM L99A-T4L uniformly [^13^C,^15^N]-labelled sample using a standard Bruker pulse sequences (hsqcctetgpsp) with coherence-selection gradients. The data was acquired with 768 and 128 (30.4 ms constant time) or 60 (15.2 constant-time) complex points in the ^1^H and ^13^C dimensions, respectively, with spectral widths of 10000 Hz and 5000 Hz. An interscan delay of 1 s was used and the data was recorded at 278K.

HMQC-type Spectra of L99A-T4L used to assess out-of-scope behaivour of FID-Net-2 and with poor shimming (Figure S5) were recorded on a 1 mM uniformly [^13^C,^15^N]-labelled samples using the pulse sequences described in Figure S3a, with τ_coup_ = 0.0 ms and 2.3 ms, on a Bruker 600 MHz Avance HD spectrometer equipped with Z-gradient triple-resonance TCI cryoprobe. The data was acquired with 768 and 200 complex points in the ^1^H and ^13^C dimensions, respectively, with spectral widths of 9009 Hz and 5000 Hz. The datasets were recorded as pseudo-3D spectra. An interscan delay of 1 s was used.

#### Three-dimensional NOESY spectra

The 3D ^1^H-^13^C-^1^H NOESY dataset of L99A-T4L (Figure 5a) used as input for FID-Net-2 were recorded on a uniformly [^13^C,^15^N]-labelled sample using the pulse sequences described in Figure S3a,d, with τ_coup_ = 0.0 ms and 2.3 ms, on a Bruker 700 MHz Avance III spectrometer equipped with Z-gradient triple-resonance TCI cryoprobe. The data was acquired with 1024, 128, and 96 complex points in ^1^H, ^1^H_NOESY_, and ^13^C_Aro_ dimensions, respectively, with spectral widths of 14280 Hz (^1^H), 5000 Hz (^13^C), and 8000 Hz (^1^H_NOESY_). Four scans were collected per increment with a recycle delay of 1 s. The mixing time was 100 ms.

The 3D ^1^H-^13^C_Aro_-^13^C_Methyl_ datasets of L99A-T4L (Figure 5c) and A39G-FF used as input for FID-Net-2 were recorded on uniformly [^13^C,^15^N]-labelled samples using the pulse sequences described in Figure S8, with τ_coup_ = 0.0 ms and 2.3 ms, on a Bruker 700 MHz Avance III spectrometer equipped with Z-gradient triple-resonance TCI cryoprobe. The data was acquired with 1024, 128, and 80 complex points in ^1^H, ^13^C_NOESY_, and ^13^C_Aro_ dimensions, respectively, with spectral widths of 14280 Hz (^1^H), 5000 Hz (^13^C_Aro_), and 5000 Hz (^1^C_Methyl_). Four scans were collected per increment with a recycle delay of 1 s. The mixing time was 120 ms for the L99A-T4L sample and 200 ms for the A39F-FF sample.

#### Longitudinal exchange (EXSY; ZZ exchange) spectra

The longitudinal aromatic ^13^C,^1^H exchange dataset of A39G-FF (Figure 6) was recorded on a 0.5 mM uniformly [^13^C,^15^N]-labelled samples using the pulse sequences described in Figure S3c, with τ_coup_ = 0.0 ms and 2.3 ms, on a Bruker 600 MHz Avance HD spectrometer equipped with Z-gradient triple-resonance TCI cryoprobe. The sample was dissolved in H_2_O buffer. An interscan delay of 1 s was used and the data was recorded at 274K. The data was acquired with 768 and 128 complex points in ^1^H and ^13^C_Aro_ dimensions, respectively, with spectral widths of 9009 Hz (^1^H), 5000 Hz (^13^C). Sixteen scans were collected per increment with a recycle delay of 1 s. The exchange delays were 25 ms, 50 ms, 100 ms, 150 ms, 200 ms, 250 ms and 300 ms.

The longitudinal aromatic ^15^N,^1^H exchange dataset of A39G-FF (Figure 6) was recorded on a 0.5 mM uniformly [^13^C,^15^N]-labelled samples using a standard pulse sequence. An interscan delay of 1 s was used and the data was recorded at 274K. The data was acquired with 1536 and 128 complex points in ^1^H and ^15^N dimensions, respectively, with spectral widths of 10000 Hz (^1^H), 2136 Hz (^13^C). Eight scans were collected per increment with a recycle delay of 1 s. The exchange delays were 25 ms (duplicate), 50 ms, 100 ms, 150 ms, 200 ms (duplicate), 250 ms, 300 ms, 400 ms, 600 ms, and 800 ms.

#### Data processing

All experimental NMR spectra were processed with NMRPIPE (*50*) or using the python libraries NMRGLUE (*51*) and NUMPY.

## Data availability

The experimental data is available from the corresponding author upon request.

## Code availability

The code for processing ^13^C-^1^H HSQC/HMQC spectra with FID-Net-2 is available from the corresponding author upon request.

## Acknowledgements

Dr Luke Nightingale is acknowledged for helpful discussions. The BBSRC (BB/R000255/1), Wellcome Trust (ref. 101569/z/13/z), and the EPSRC are acknowledged for supporting the NMR facility at University College London. Access to ultra-high field NMR spectrometers was supported by the Francis Crick Institute through provision of access to the MRC Biomedical NMR Centre. The Francis Crick Institute receives its core funding from Cancer Research UK (FC001029), the UK Medical Research Council (FC001029), and the Wellcome Trust (FC001029). This study made use of NMRbox: National Center for Biomolecular NMR Data Processing and Analysis (*48*), a Biomedical Technology Research Resource (BTRR), which is supported by NIH grant P41GM111135 (NIGMS). Some computational aspects of this work were supported by the Francis Crick Institute (DFH) through provision of access to the Scientific Computing STP and the Crick data Analysis and Management Platform (CAMP). The Francis Crick Institute (CAMP) receives its core funding from Cancer Research UK (FC010233), the UK Medical Research Council (FC010233), and the Wellcome Trust (FC010233). PV acknowledges intramural funding from TIFR Hyderabad (DAE, Government of India, Project No. RTI 4007). For the purpose of open access, the author has applied a Creative Commons Attribution (CC BY) licence to any Author Accepted Manuscript version arising. This research is supported by the UKRI and EPSRC (EP/X036782/1).

## Author information

V.K.S., G.K., P.V. and D.F.H designed the research; D.F.H. designed and trained all the DNNs; V.K.S. produced all isotope labelled samples; V.K.S., P.V. and D.F.H. performed and analysed NMR experiments; V.K.S assigned the chemical shifts. All of the authors analysed the data, discussed the results and wrote the paper.

## References

1. T. R. Alderson, L. E. Kay, NMR spectroscopy captures the essential role of dynamics in regulating biomolecular function. Cell. 184, 577–595 (2021).

2. A. G. Palmer, Enzyme Dynamics from NMR Spectroscopy. Acc. Chem. Res. 48, 457–465 (2015).

3. T. Xie, T. Saleh, P. Rossi, C. G. Kalodimos, Conformational states dynamically populated by a kinase determine its function. Science. 370, eabc2754 (2020).

4. L. Mariño Pérez, F. S. Ielasi, L. M. Bessa, D. Maurin, J. Kragelj, M. Blackledge, N. Salvi, G. Bouvignies, A. Palencia, M. R. Jensen, Visualizing protein breathing motions associated with aromatic ring flipping. Nature. 602, 695–700 (2022).

5. J. B. Stiller, R. Otten, D. Häussinger, P. S. Rieder, D. L. Theobald, D. Kern, Structure determination of high-energy states in a dynamic protein ensemble. Nature. 603, 528–535 (2022).

6. K. Madhurima, B. Nandi, S. Munshi, A. N. Naganathan, A. Sekhar, Functional regulation of an intrinsically disordered protein via a conformationally excited state. Sci. Adv. 9, eadh4591 (2023).

7. V. K. Shukla, L. Siemons, D. F. Hansen, Intrinsic structural dynamics dictate enzymatic activity and inhibition. Proc. Natl. Acad. Sci. 120, e2310910120 (2023).

8. H. W. Mackenzie, D. F. Hansen, Arginine Side-Chain Hydrogen Exchange: Quantifying Arginine Side-Chain Interactions in Solution. ChemPhysChem. 20, 252–259 (2019).

9. A. Ceccon, V. Tugarinov, F. Torricella, G. M. Clore, Quantitative NMR analysis of the kinetics of prenucleation oligomerization and aggregation of pathogenic huntingtin exon-1 protein. Proc. Natl. Acad. Sci. 119, e2207690119 (2022).

10. S. Guseva, V. Schnapka, W. Adamski, D. Maurin, R. W. H. Ruigrok, N. Salvi, M. Blackledge, Liquid–Liquid Phase Separation Modifies the Dynamic Properties of Intrinsically Disordered Proteins. J. Am. Chem. Soc. 145, 10548–10563 (2023).

11. S. Vahidi, Z. A. Ripstein, J. B. Juravsky, E. Rennella, A. L. Goldberg, A. K. Mittermaier, J. L. Rubinstein, L. E. Kay, An allosteric switch regulates Mycobacterium tuberculosis ClpP1P2 protease function as established by cryo-EM and methyl-TROSY NMR. Proc. Natl. Acad. Sci. 117, 5895–5906 (2020).

12. V. K. Shukla, G. T. Heller, D. F. Hansen, Biomolecular NMR spectroscopy in the era of artificial intelligence. Structure. 31, 1360–1374 (2023).

13. J. Jumper, R. Evans, A. Pritzel, T. Green, M. Figurnov, O. Ronneberger, K. Tunyasuvunakool, R. Bates, A. Žídek, A. Potapenko, A. Bridgland, C. Meyer, S. A. A. Kohl, A. J. Ballard, A. Cowie, B. Romera-Paredes, S. Nikolov, R. Jain, J. Adler, T. Back, S. Petersen, D. Reiman, E. Clancy, M. Zielinski, M. Steinegger, M. Pacholska, T. Berghammer, S. Bodenstein, D. Silver, O. Vinyals, A. W. Senior, K. Kavukcuoglu, P. Kohli, D. Hassabis, Highly accurate protein structure prediction with AlphaFold. Nature. 596, 583–589 (2021).

14. K. H. Gardner, L. E. Kay, The use of ^2^H, ^13^C, ^15^N multidimensional NMR to study the structure and dynamics of proteins. Annu. Rev. Biophys. Biomol. Struct. 27, 357–406 (1998).

15. K. Pervushin, R. Riek, G. Wider, K. Wüthrich, Attenuated *T*_2_ relaxation by mutual cancellation of dipole-dipole coupling and chemical shift anisotropy indicates an avenue to NMR structures of very large biological macromolecules in solution. Proc. Natl. Acad. Sci. USA. 94, 12366–12371 (1997).

16. V. Tugarinov, P. M. Hwang, J. E. Ollerenshaw, L. E. Kay, Cross-Correlated Relaxation Enhanced ^1^H−^13^C NMR Spectroscopy of Methyl Groups in Very High Molecular Weight Proteins and Protein Complexes. J. Am. Chem. Soc. 125, 10420–10428 (2003).

17. N. D. Werbeck, J. Kirkpatrick, D. F. Hansen, Probing arginine side-chains and their dynamics with carbon-detected NMR spectroscopy: application to the 42 kDa human histone deacetylase 8 at high pH. Angew. Chem. Int. Ed. Engl. 52, 3145–3147 (2013).

18. R. B. Pritchard, D. F. Hansen, Characterising side chains in large proteins by protonless 13C-detected NMR spectroscopy. Nat. Commun. 10, 1747 (2019).

19. A. Esadze, C. Chen, L. Zandarashvili, S. Roy, B. M. Pettitt, J. Iwahara, Changes in conformational dynamics of basic side chains upon protein–DNA association. Nucleic Acids Res. 44, 6961–6970 (2016).

20. K. A. Stafford, F. Ferrage, J.-H. Cho, A. G. Palmer, Side Chain Dynamics of Carboxyl and Carbonyl Groups in the Catalytic Function of Escherichia coli Ribonuclease H. J. Am. Chem. Soc. 135, 18024–18027 (2013).

21. J. Santoro, G. C. King, A constant-time 2D overbodenhausen experiment for inverse correlation of isotopically enriched species. J. Magn. Reson. 97, 202–207 (1992).

22. G. W. Vuister, A. Bax, Resolution enhancement and spectral editing of uniformly ^13^C-enriched proteins by homonuclear broadband 13C decoupling. J. Magn. Reson. 98, 428–435 (1992).

23. K. Teilum, U. Brath, P. Lundström, M. Akke, Biosynthetic ^13^C Labeling of Aromatic Side Chains in Proteins for NMR Relaxation Measurements. J. Am. Chem. Soc. 128, 2506–2507 (2006).

24. M. Akke, U. Weininger, NMR Studies of Aromatic Ring Flips to Probe Conformational Fluctuations in Proteins. J. Phys. Chem. B. 127, 591–599 (2023).

25. U. Weininger, Optimal Isotope Labeling of Aromatic Amino Acid Side Chains for NMR Studies of Protein Dynamics. Methods Enzymol. 614, 67–86 (2019).

26. B. M. Young, P. Rossi, P. J. Slavish, Y. Cui, M. Sowaileh, J. Das, C. G. Kalodimos, Z. Rankovic, Synthesis of Isotopically Labeled, Spin-Isolated Tyrosine and Phenylalanine for Protein NMR Applications. Org. Lett. 23, 6288–6292 (2021).

27. Y. LeCun, Y. Bengio, G. Hinton, Deep learning. Nature. 521, 436–444 (2015).

28. M. Baek, F. DiMaio, I. Anishchenko, J. Dauparas, S. Ovchinnikov, G. R. Lee, J. Wang, Q. Cong, L. N. Kinch, R. D. Schaeffer, C. Millán, H. Park, C. Adams, C. R. Glassman, A. DeGiovanni, J. H. Pereira, A. V. Rodrigues, A. A. van Dijk, A. C. Ebrecht, D. J. Opperman, T. Sagmeister, C. Buhlheller, T. Pavkov-Keller, M. K. Rathinaswamy, U. Dalwadi, C. K. Yip, J. E. Burke, K. C. Garcia, N. V. Grishin, P. D. Adams, R. J. Read, D. Baker, Accurate prediction of protein structures and interactions using a three-track neural network. Science. 373, 871–876 (2021).

29. S. G. Worswick, J. A. Spencer, G. Jeschke, I. Kuprov, Deep neural network processing of DEER data. Sci. Adv. 4, eaat5218 (2018).

30. X. Qu, Y. Huang, H. Lu, T. Qiu, D. Guo, T. Agback, V. Orekhov, Z. Chen, Accelerated Nuclear Magnetic Resonance Spectroscopy with Deep Learning. Angew. Chemie. 132, 10383–10386 (2020).

31. G. Karunanithy, D. F. Hansen, FID-Net: A versatile deep neural network architecture for NMR spectral reconstruction and virtual decoupling. J. Biomol. NMR. 75, 179–191 (2021).

32. G. Karunanithy, H. W. Mackenzie, D. F. Hansen, Virtual Homonuclear Decoupling in Direct Detection Nuclear Magnetic Resonance Experiments Using Deep Neural Networks. J. Am. Chem. Soc. 143, 16935–16942 (2021).

33. D.-W. Li, A. L. Hansen, C. Yuan, L. Bruschweiler-Li, R. Brüschweiler, DEEP picker is a deep neural network for accurate deconvolution of complex two-dimensional NMR spectra. Nat. Commun. 12, 5229 (2021).

34. D. F. Hansen, Using Deep Neural Networks to Reconstruct Non-uniformly Sampled NMR Spectra. J. Biomol. NMR. 73, 577–585 (2019).

35. G. Karunanithy, V. K. Shukla, D. F. Hansen, Solution State Methyl NMR Spectroscopy of Large Non-Deuterated Proteins Enabled by Deep Neural Networks. bioRxiv (2023), doi:10.1101/2023.09.15.557823.

36. A. E. Eriksson, W. A. Baase, J. A. Wozniak, B. W. Matthews, A cavity-containing mutant of T4 lysozyme is stabilized by buried benzene. Nature. 355, 371–373 (1992).

37. Y. Pustovalova, F. Delaglio, D. L. Craft, H. Arthanari, A. Bax, M. Billeter, M. J. Bostock, H. Dashti, D. F. Hansen, S. G. Hyberts, B. A. Johnson, K. Kazimierczuk, H. Lu, M. Maciejewski, T. M. Miljenović, M. Mobli, D. Nietlispach, V. Orekhov, R. Powers, X. Qu, S. A. Robson, D. Rovnyak, G. Wagner, J. Ying, M. Zambrello, J. C. Hoch, D. L. Donoho, A. D. Schuyler, NUScon: a community-driven platform for quantitative evaluation of nonuniform sampling in NMR. Magn. Reson. 2, 843–861 (2021).

38. G. Bouvignies, P. Vallurupalli, D. F. Hansen, B. E. Correia, O. Lange, A. Bah, R. M. Vernon, F. W. Dahlquist, D. Baker, L. E. Kay, Solution structure of a minor and transiently formed state of a T4 lysozyme mutant. Nature. 477, 111–117 (2011).

39. L. Liu, W. A. Baase, B. W. Matthews, Halogenated Benzenes Bound within a Non-polar Cavity in T4 Lysozyme Provide Examples of I⋯S and I⋯Se Halogen-bonding. J. Mol. Biol. 385, 595– 605 (2009).

40. M. Tollinger, N. R. Skrynnikov, F. A. A. Mulder, J. D. Forman-Kay, L. E. Kay, Slow dynamics in folded and unfolded states of an SH3 domain. J. Am. Chem. Soc. 123, 11341–11352 (2001).

41. V. P. Tiwari, Y. Toyama, D. De, L. E. Kay, P. Vallurupalli, The A39G FF domain folds on a volcano-shaped free energy surface via separate pathways. Proc. Natl. Acad. Sci. 118, e2115113118 (2021).

42. H. M. McConnell, Reaction rates by nuclear magnetic resonance. J. Chem. Phys. 28, 430–431 (1958).

43. A. Boeszoermenyi, S. Chhabra, A. Dubey, D. L. Radeva, N. T. Burdzhiev, C. D. Chanev, O. I. Petrov, V. M. Gelev, M. Zhang, C. Anklin, H. Kovacs, G. Wagner, I. Kuprov, K. Takeuchi, H. Arthanari, Aromatic ^19^F-^13^C TROSY: a background-free approach to probe biomolecular structure, function, and dynamics. Nat. Methods. 16, 333–340 (2019).

44. L.-P. Picard, R. S. Prosser, Advances in the study of GPCRs by ^19^F NMR. Curr. Opin. Struct. Biol. 69, 169–176 (2021).

45. M. Abadi, A. Agarwal, P. Barham, E. Brevdo, Z. Chen, C. Citro, G. S. Corrado, A. Davis, J. Dean, M. Devin, S. Ghemawat, I. Goodfellow, A. Harp, G. Irving, M. Isard, R. Jozefowicz, Y. Jia, L. Kaiser, M. Kudlur, J. Levenberg, D. Mané, M. Schuster, R. Monga, S. Moore, D. Murray, C. Olah, J. Shlens, B. Steiner, I. Sutskever, K. Talwar, P. Tucker, V. Vanhoucke, V. Vasudevan, F. Viégas, O. Vinyals, P. Warden, M. Wattenberg, M. Wicke, Y. Yu, X. Zheng, TensorFlow: Large-scale machine learning on heterogeneous systems (2015), (available at www.tensorflow.org).

46. F. and others Chollet, Keras (2015), (available at https://keras.io).

47. D. P. Kingma, J. Ba, Adam: A Method for Stochastic Optimization (2014), doi:1412.6980.

48. M. W. Maciejewski, A. D. Schuyler, M. R. Gryk, I. I. Moraru, P. R. Romero, E. L. Ulrich, H. R. Eghbalnia, M. Livny, F. Delaglio, J. C. Hoch, NMRbox: A Resource for Biomolecular NMR Computation. Biophys. J. 112, 1529–1534 (2017).

49. K. H. Gardner, X. Zhang, K. Gehring, L. E. Kay, Solution NMR Studies of a 42 KDa Escherichia C oli Maltose Binding Protein/β-Cyclodextrin Complex: Chemical Shift Assignments and Analysis. J. Am. Chem. Soc. 120, 11738–11748 (1998).

50. F. Delaglio, S. Grzesiek, G. W. Vuister, G. Zhu, J. Pfeifer, A. Bax, Nmrpipe - a Multidimensional Spectral Processing System Based on Unix Pipes. J. Biomol. Nmr. 6, 277– 293 (1995).

51. J. J. Helmus, C. P. Jaroniec, Nmrglue: an open source Python package for the analysis of multidimensional NMR data. J. Biomol. NMR. 55, 355–367 (2013).

