## Supplementary Information for "Characterising Aromatic Side Chains in Proteins through the Synergistic Development of NMR Experiments and Deep Neural Networks"

### Supporting Tables:

**Table S1. Parameters used for Training**

| **Parameter** | |
|---|---|
| Number of cross-peaks | 40 – 200 (uniform distribution) |
| Intensity | $\mathcal{N}(1, 0.5)$[a)] |
| Noise | $\mathcal{N}(1.2, 0.12)$ |
| $^{1}$H SW (Hz) | 2000 – 5000 |
| $^{1}$H number of complex points | 128 – 256 |
| $R_2(^{1}H)$ ($s^{-1}$) | $\lvert \mathcal{N}(50, 20) \rvert$ |
| Phase($^{1}$H) | $\mathcal{N}(0, 10)$ |
| $J_{HH}$ (Hz) | $\mathcal{N}(8, 2)$ [10% set to zero] |
| $J_{HH}$ (Hz) | $\mathcal{N}(4, 2)$ [50% set to zero] |
| $^{13}$C SW (Hz) | 4000 – 6650 |
| $^{13}$C Acquisition time (s) | 0.030 – 0.050 (uniform distribution) |
| $^{13}$C number of complex points | 96 – 200 (uniform distribution) |
| $R_2(^{13}C)$ ($s^{-1}$) | $\lvert \mathcal{N}(0, 1) \rvert$ |
| Phase($^{13}$C) | $\mathcal{N}(0, 10)$ |
| $J_{CC}$ (Hz) | $\mathcal{N}(63, 10)$ [20% set to zero] |
| $J_{CC}$ (Hz) | $\mathcal{N}(63, 10)$ [20% set to zero] |

a) $\mathcal{N}(\mu, \sigma)$ is a normal distribution with mean $\mu$ and standard deviation $\sigma$.

### Supporting Figures:

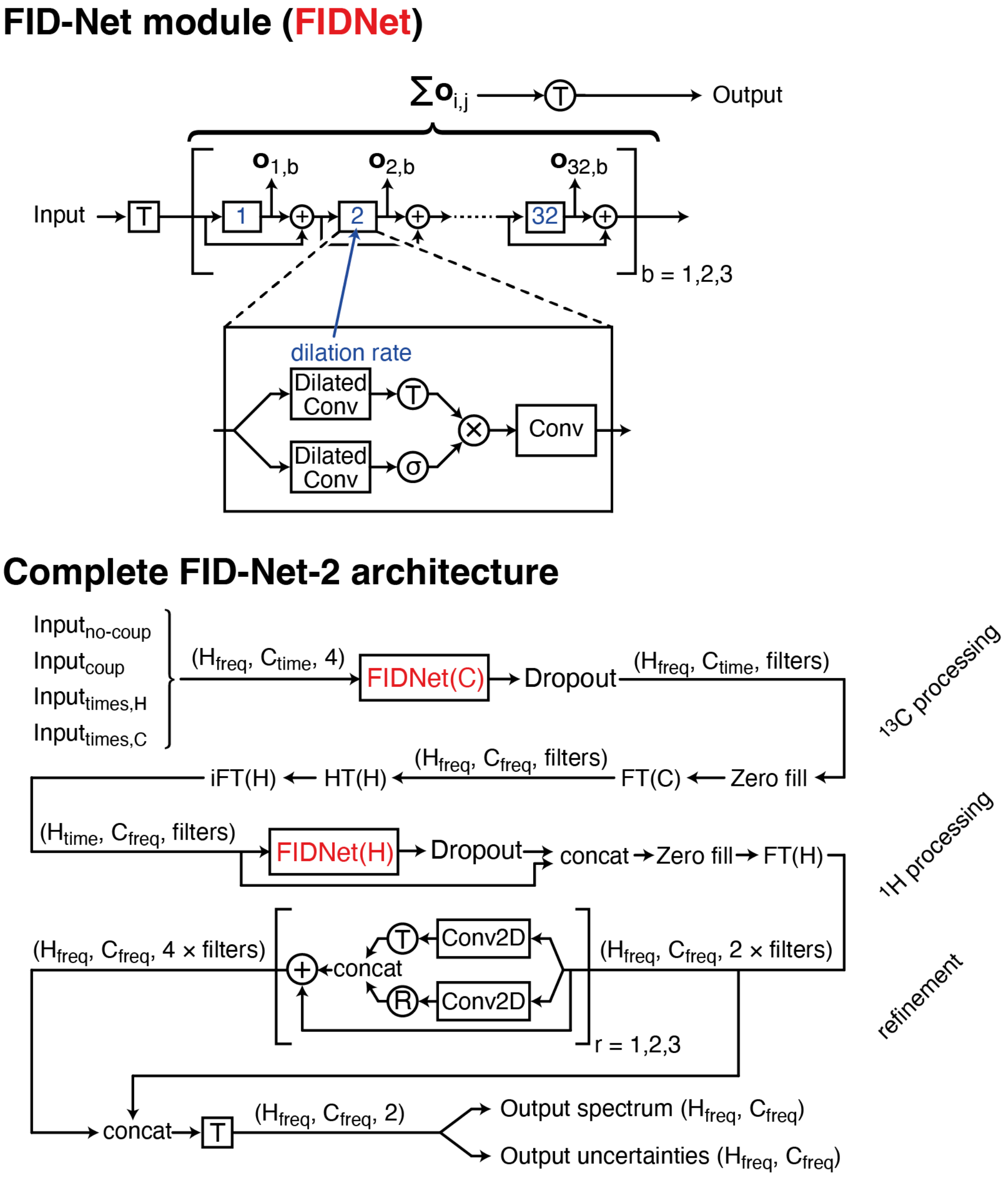

**Figure S1: Summary of the FID-Net-2 Network architecture.** (**a**) the FID-Net block, which is very similar to the FID-Net architecture published previously (*1*). (**b**) The FID-Net-2 architecture, which consists of mainly three parts, (*i*) a transformation of the $^{13}$C dimension with an FID-Net module, (*ii*) a transformation of the $^{1}$H dimension with an FID-Net module, and (*iii*) a refinement. Circles denote elementwise transformations: R: rectified linear unit, T: tanh($x$) operation, σ: sigmoidal $1/(\exp(-x)+1)$ operation, +: summation, ×: multiplication. These elementwise operations do not include weights to be optimised. Rectangles denote layers with trainable weights: Conv: Convolutional layer, Conv2D: two-dimensional convolutional layer, T: a dense linear layer with tanh($x$) activation function. The dropout rate used for the Dropout layers was 10% and was only applied during training. FT donate a Fourier transform, iFT is an inverse Fourier transform, and HT is a Hilbert transformation.

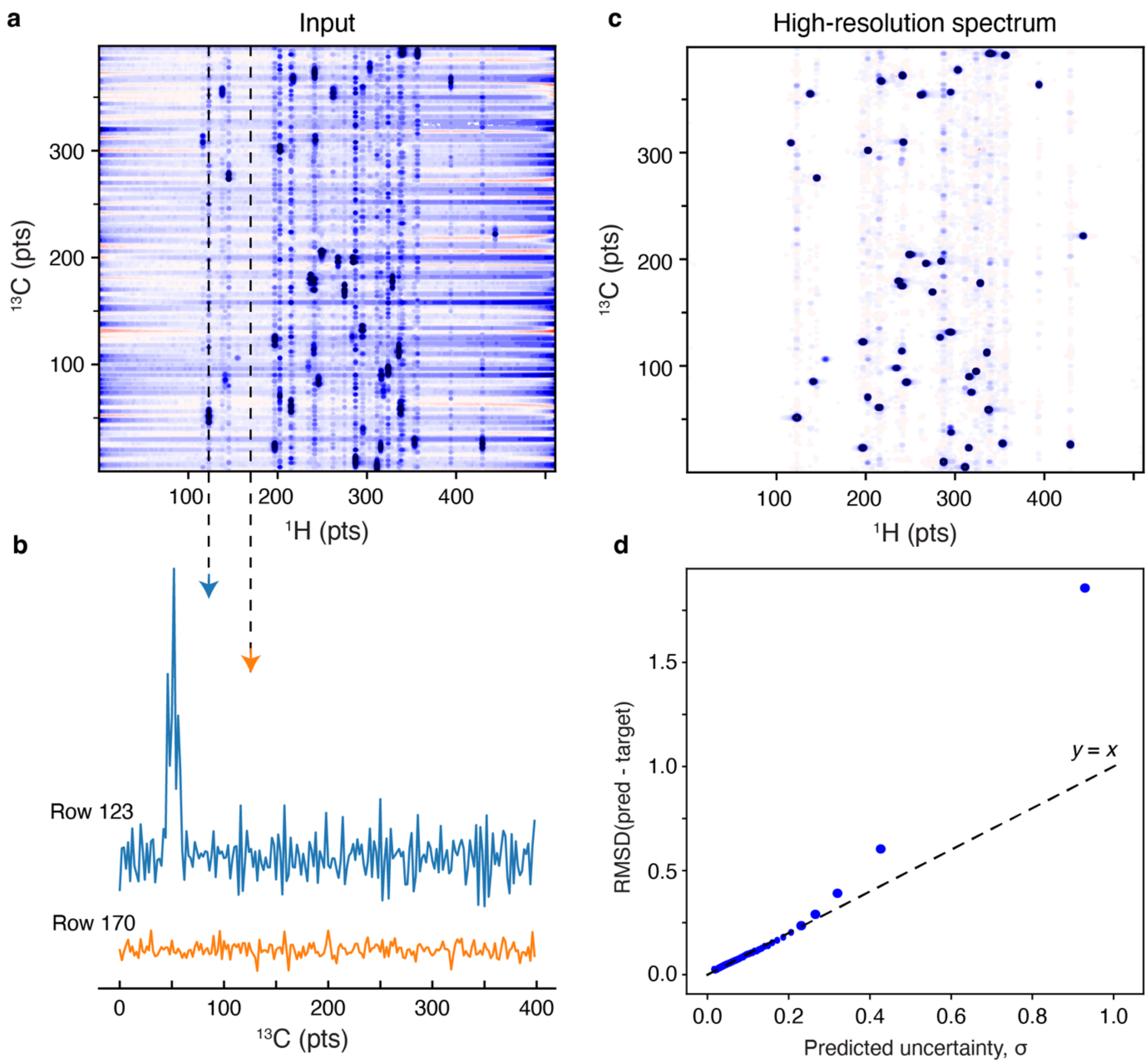

**Figure S2: Assessing the performance of FID-Net2 in the presence of $t_1$-noise using synthetic data.** (**a**) A representative simulated spectrum, corresponding to a 20 kDa protein at 298K. On average 50 peaks are present with transverse relaxation rates of 45 $s^{-1}$ in both the $^{1}$H and $^{13}$C dimension. The $t_1$-noise was simulated by adding Gaussian noise to the $t_1$ evolution time, in this case, $t_1(n) = n/\mathrm{SW} \times \mathcal{N}(1, 0.005)$, where SW is the sweep width in Hz and $\mathcal{N}(\mu, \sigma)$ is a normal distribution with mean $\mu$ and standard deviation $\sigma$. (**b**) Slices from the spectrum in **a**, at a position where a cross-peak is present (row 123) and at a position where only noise is expected (row 170). (**c**) Spectrum obtained after processing with the FID-Net2. Although the DNN was not trained on spectra containing $t_1$ noise, it is seen that the processing is robust, albeit the result not being as good as processing spectra without $t_1$ noise. (**d**) Predicted uncertainties *v.s.* calculated RMSD for 10 random spectra over 200 bins. The uncertainties are underestimated when substantial $t_1$ noise is present, however, the processing with FID-Net-2 is general robust.

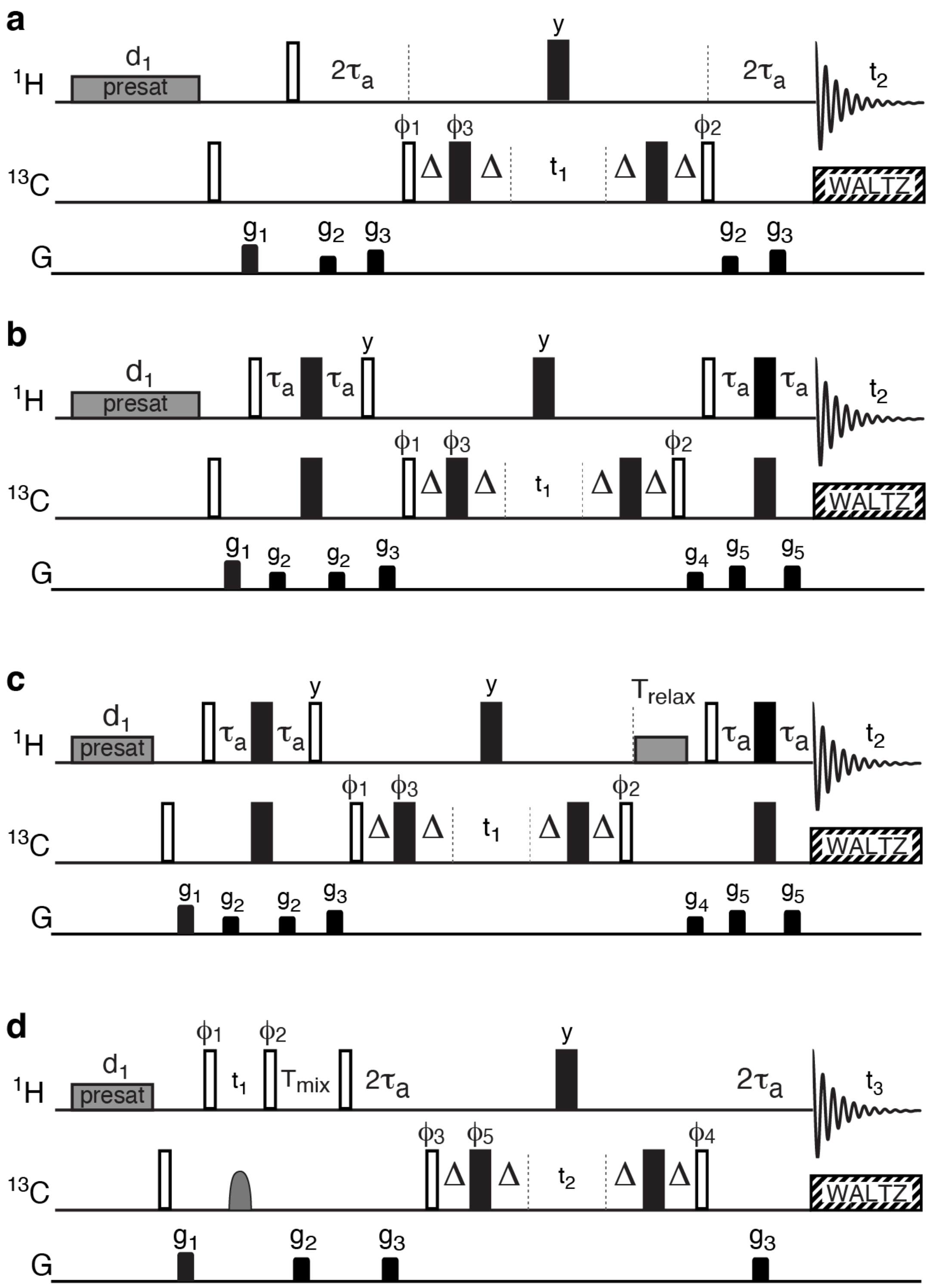

**Figure S3: Pulse sequences used to generate $^{13}C$-$^{1}H$ correlation spectra for FID-Net-2 processing.** Common to all sequences, (**a-d**), is that black bars represent 180° pulses, whereas open bars represent 90° non-selective pulses, that are all applied at the highest available field. The $^{1}H$ carrier is placed on the $H_2O$ signal (~ 4.77 ppm) and the $^{13}C$ carrier placed in the middle of the aromatic region (122 ppm relative to TMS). The delays used are $\tau_a = 1/(4\ J_{CH}(\text{aro})\ ) = 1.4$ ms, whereas two planes are recorded one with $\Delta = 0.0$ and another with $\Delta = 2.3$ ms. Pre-saturation of the $H_2O$ solvent signal is achieved with a 35 Hz pulse during the recovery delay $d_1$. The phase cycle for (**a-c**) is $\phi_1$: *x*, *-x*; $\phi_2$: 4(*x*), 4(*-x*); $\phi_3$: *x*, *-x*, *y*, *-y*, *-x*, *x*, *-y*, *y*; $\phi_{rec}$: *x*, *-x*, *-x*, *x*, *-x*, *x*, *x*, *-x*. Frequency discrimination in $t_1$ is obtained by States-TPPI

(*3*) of the phase $\phi_1$ (States) and $\phi_2$ (TPPI). (**a**) HMQC-type 2D $^{13}$C-$^1$H correlation map. Gradients of 0.5 ms are represented by black rectangles and applied with strength of $g_1$: 3.4 G/cm, $g_2$: 0.8 G/cm, $g_3$: 1.3 G/cm. (**b**) HSQC-type 2D $^{13}$C-$^1$H correlation map. Gradients of 0.5 ms are represented by black rectangles and applied with strength of $g_1$: 3.4 G/cm, $g_2$: 0.8 G/cm, $g_3$: 4.5 G/cm, $g_4$: 6.1 G/cm, $g_5$: 1.2 G/cm. (**c**) HSQC longitudinal exchange experiment. Gradients of 0.5 ms are represented by black rectangles and applied with strength of $g_1$: 3.4 G/cm, $g_2$: 0.8 G/cm, $g_3$: 4.5 G/cm, $g_4$: 6.1 G/cm, $g_5$: 1.2 G/cm. (**d**) $^1$H-$^{13}$C-$^1$H NOESY experiment. The gray shaped pulse is an adiabatic inversion pulse, shape Crp80,0.5,20.1, which is applied for 500 μs and leading to inversion of a ±20 kHz frequency range. The phase cycle is $\phi_1$: *x*; $\phi_2$: 2(*x*), 2(-*x*); $\phi_3$: *x*, -*x*; $\phi_1$: *x*; $\phi_5$: *x*, -*x*, *y*, -*y*, -*x*, *x*, -*y*, *y*; $\phi_{rec}$: *x*, -*x*, -*x*, *x*, -*x*, *x*, *x*, -*x*. Frequency discrimination in $t_1$ is obtained by States-TPPI (*3*) of the phase $\phi_1$ (States) and $\phi_2$ (TPPI). Frequency discrimination in $t_2$ is obtained by States-TPPI of the phase $\phi_3$ (States) and $\phi_4$ (TPPI). Gradients of 0.2 ms are represented by black rectangles and applied with strength of $g_1$: 3.4 G/cm, $g_2$: 0.8 G/cm, $g_3$: 4.0 G/cm.

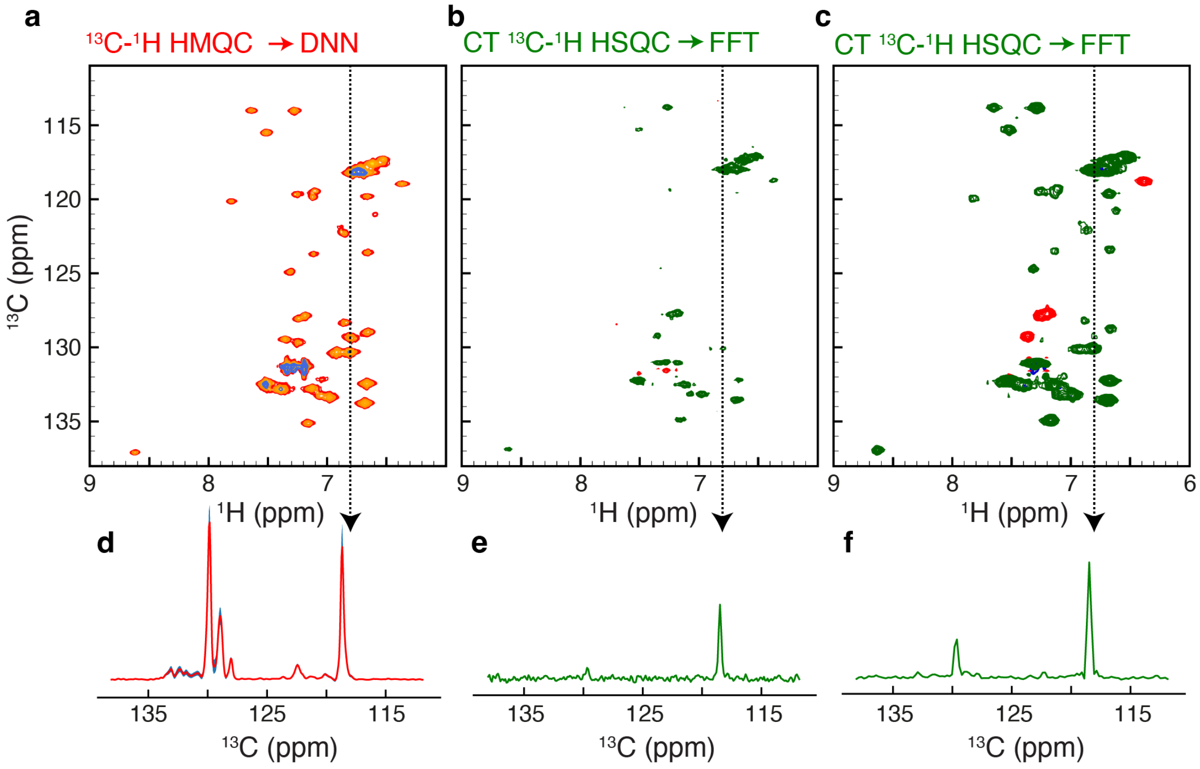

**Figure S4: Comparison between Fid-Net-2 transformed spectra and constant-time HSQC $^{13}$C-$^{1}$H correlation maps.** (**a**) Fid-Net-2 transformed $^{13}$C-$^{1}$H spectrum (red-yellow) along with the uncertainties (blue) of L99A-T4L (~1 mM) recorded with 8 scans. (**b**) Constant-time $^{13}$C-$^{1}$H HSQC recorded with a constant-time delay of 30.4 ms and recorded with 16 scans (**c**) Constant-time $^{13}$C-$^{1}$H HSQC recorded with a constant-time delay of 15.2 ms and 16 scans. Cross-peaks with negative intensity (red) stem from $^{13}$C nuclei only scalar coupled to one adjacent $^{13}$C. All spectra were recorded at a static magnetic field of 16.4 T (700 MHz) and at a temperature of 278 K.

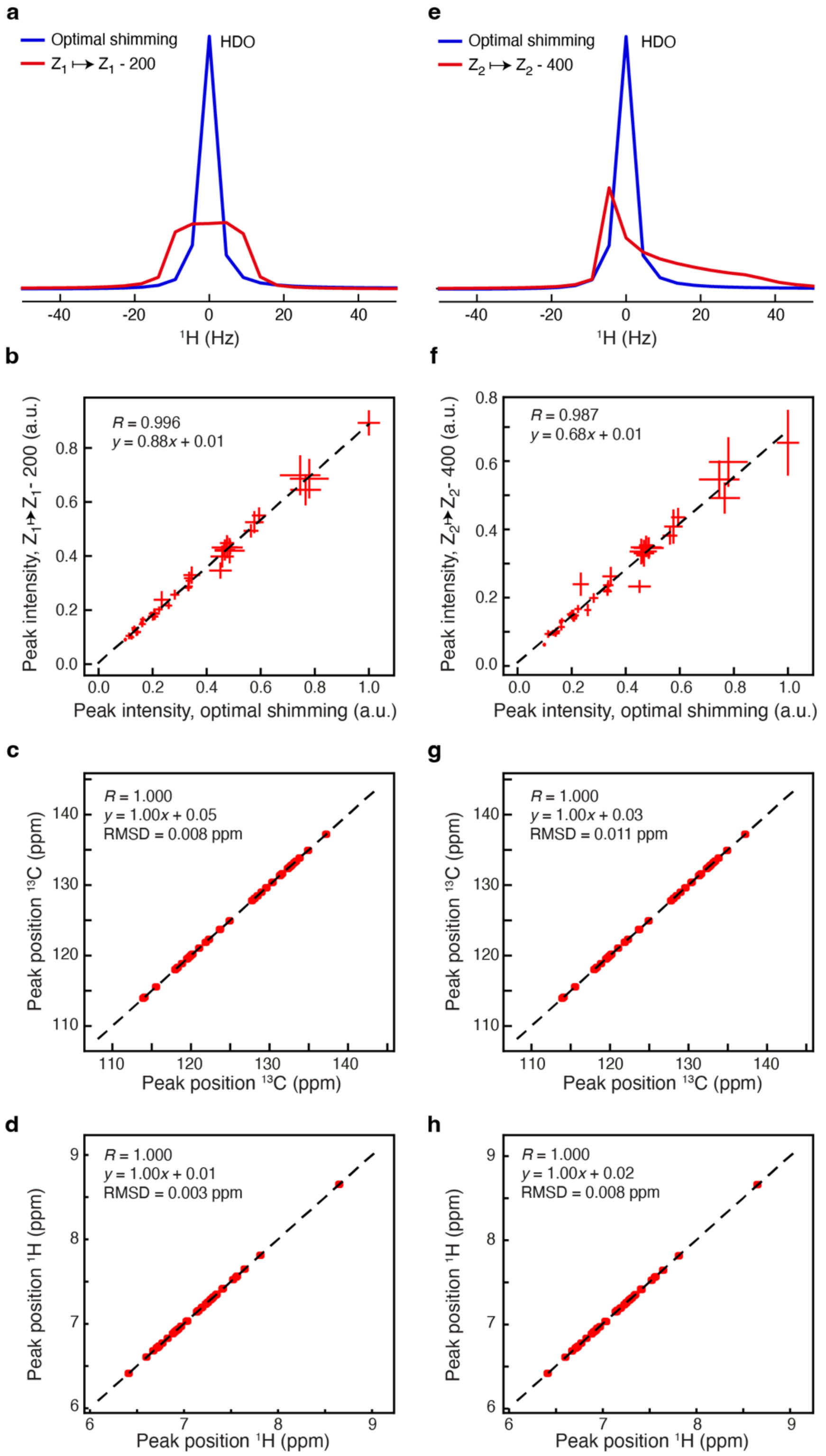

**Figure S5: Assessing the performance of Fid-Net-2 on datasets collected with poor shims.** The assessment is carried out on L99A-T4L on data recorded at 600 MHz and a temperature of 298K. (**a,e**) One-dimensional spectra with optimal and poor shimming, when the $Z_1$ (**a**) or $Z_2$ (**e**) shims have been offset. (**b,c,d**) Assessment of peak-intensities (**b**), $^{13}$C peak positions (**c**), and $^{1}$H peak positions (**d**) when the $Z_1$ shim has been offset. (**f,g,h**) Assessment of peak-intensities (**f**), $^{13}$C peak positions (**g**), and $^{1}$H peak positions (**h**) when the $Z_2$ shim has been offset. It is important to note that the DNN was only trained on optimally shimmed synthetic spectra and this assessment therefore represents an application to data that are substantially outside the training dataset.

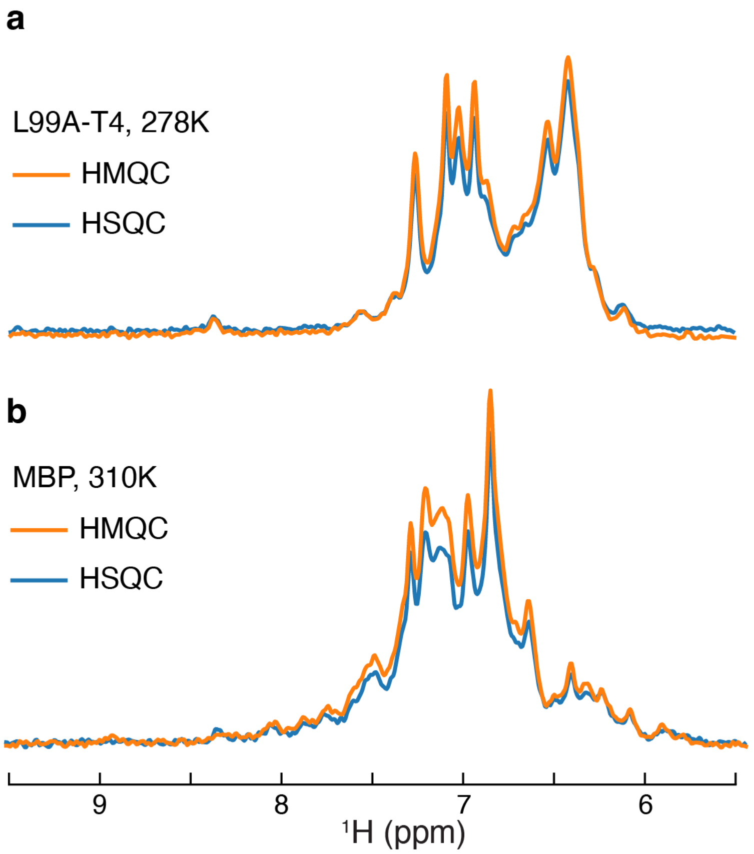

**Figure S6: Sensitivity of $^{13}$C-$^{1}$H HSQC versus $^{13}$C-$^{1}$H HMQC. (a)** Comparison of a one-dimensional $^{1}$H spectra, $^{13}$C($t_1$) = 0 s, of L99A-T4L recorded at 700 MHz, at 278K and using a HSQC-type (blue) and HMQC-type (orange). (**b)** Comparison of a one-dimensional $^{1}$H spectra, $^{13}$C($t_1$) = 0 s, of MBP recorded at 700 MHz, at 310K and using a HSQC-type (blue) and HMQC-type (orange). In both cases, the HMQC-type spectra are slightly more sensitive.

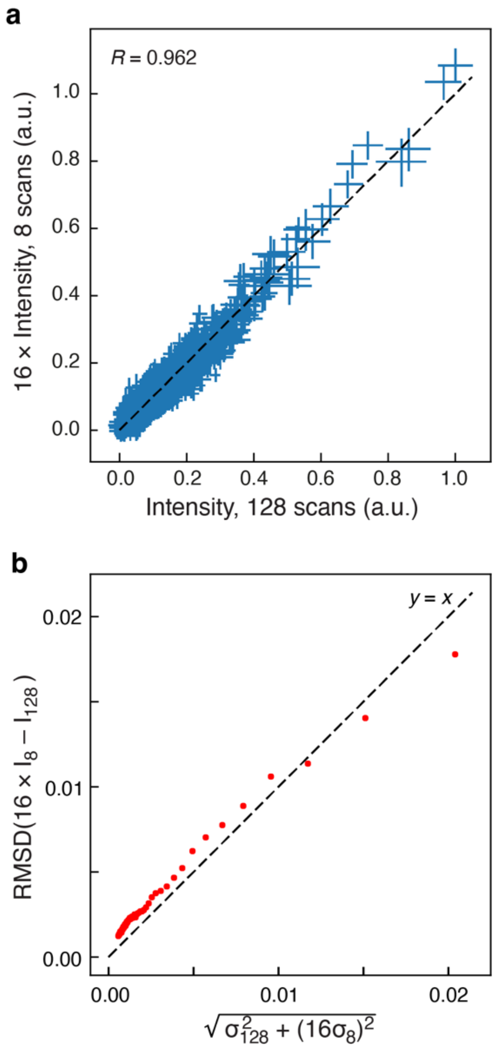

**Figure S7: Comparing the performance of Fid-Net-2 on datasets with differing signal to noise.** Two spectra of MBP (Figure 5) were recorded with 8 scans and 128 scans. Thus, the two spectra were recorded for 1.2 h and 20h, respectively. Subsequently the two spectra were processed with FID-Net-2 network, and the results compared point-by-point between the two spectra. (**a**) Comparison of all spectral values over the 512 pts ($^{1}$H) × 400 pts ($^{13}$C). Vertical and horizontal bars represent the uncertainties predicted by the DNN and *R* is the Pearson coefficient of correlation. (**b**) Comparison of differences between the two spectra, RMSD, and the estimated uncertainties. It is seen that the DNN excellently predicts reliable uncertainties, even in this case where the two spectra are obtained with 4 times different signal-to-noise. The uncertainty comparison is calculated over 200 linearly spaced bins.

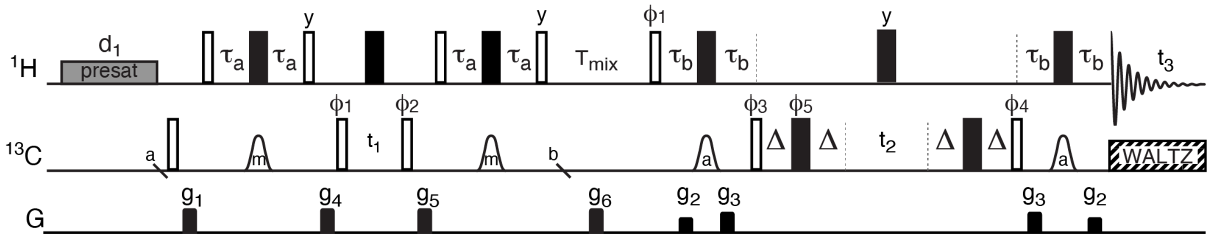

**Figure S8: Pulse sequences to obtain methyl ($^{13}CH_3$) to aromatic $^{13}C$-$^1H$ through-space NOEs.** Black bars represent 180° while open bars represent 90° non-selective pulses that are applied at the highest available field. The pre-saturation $^1H$ pulse was applied on the $H_2O$ solvent signal using a field of 35 Hz. Open bell-shaped pulses represent frequency-selective 180° pulses using a Re-BURP shape (*2*), shaped pulses with annotation 'm' are centred at the $^{13}C$ methyl region (16 ppm), whereas shaped annotated with 'a' are centred at the aromatic region (122 ppm). The length of the shaped pulses were 0.95 ms (700 MHz). The $^1H$ carrier was placed on the $H_2O$ signal, whereas the $^{13}C$ carrier was at 16 ppm (methyl region) between *a* and *b*, and otherwise at 122 ppm (aromatic region). The following delays are used: $\tau_a = 1/(4\ J_{CH}(\text{methyl})) = 2$ ms, $\tau_b = 1/(4\ J_{CH}(\text{aro})) = 1.4$ ms, $\Delta = 0.0$ and 2.3 ms, $T_{mix}$ is the NOESY mixing time. Gradients of 0.2 ms are represented by black rectangles and applied with strength of $g_1$: 3.4 G/cm, $g_2$: 0.8 G/cm, $g_3$: 4.4 G/cm, $g_4$: 3.8 G/cm, $g_5$: 6.3 G/cm, $g_6$: 2.2 G/cm. The phase cycle is $\phi_1$: *x*; $\phi_2$: 2(*x*), 2(*-x*); $\phi_3$: *x*, *-x*; $\phi_4$: *x*; $\phi_5$: *x*, *-x*, *y*, *-y*, *-x*, *x*, *-y*, *y*; $\phi_{rec}$: *x*, *-x*, *-x*, *x*, *-x*, *x*, *x*, *-x*. Frequency discrimination in $t_1$ is obtained by States-TPPI (*3*) of the phase $\phi_1$ (States) and $\phi_2$ (TPPI). Frequency discrimination in $t_2$ is obtained by States-TPPI of the phase $\phi_3$ (States) and $\phi_4$ (TPPI).

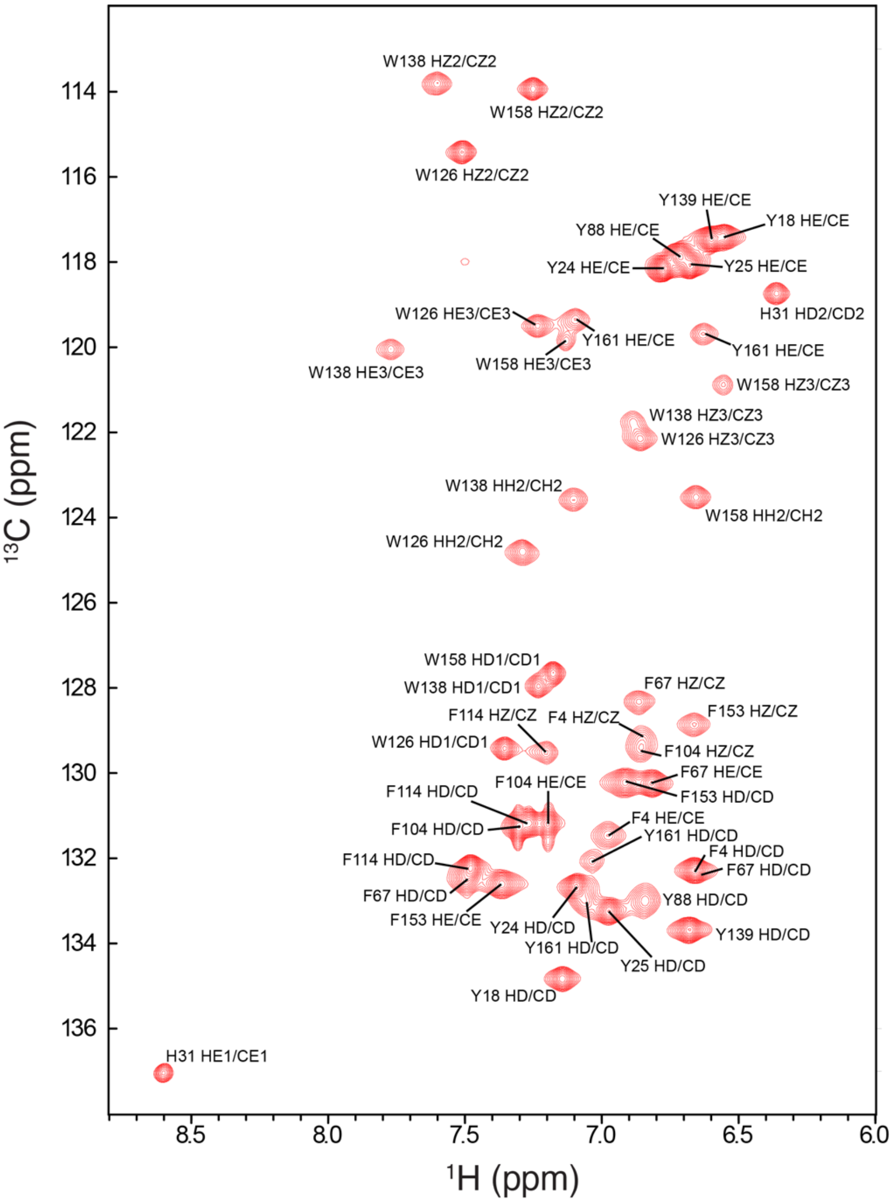

**Figure S9: Assignment of the FID-Net-2 processed aromatic $^{13}C$-$^{1}H$ HSQC spectrum of L99A-T4L (700 MHz; 298K)**. Chemical shift assignments were obtained using the $^{1}H$-$^{13}C$-$^{1}H$ NOESY-HSQC and $^{13}C$-$^{13}C$-$^{1}H$ HSQC-NOESY-HSQC spectra described in Fig. 4.

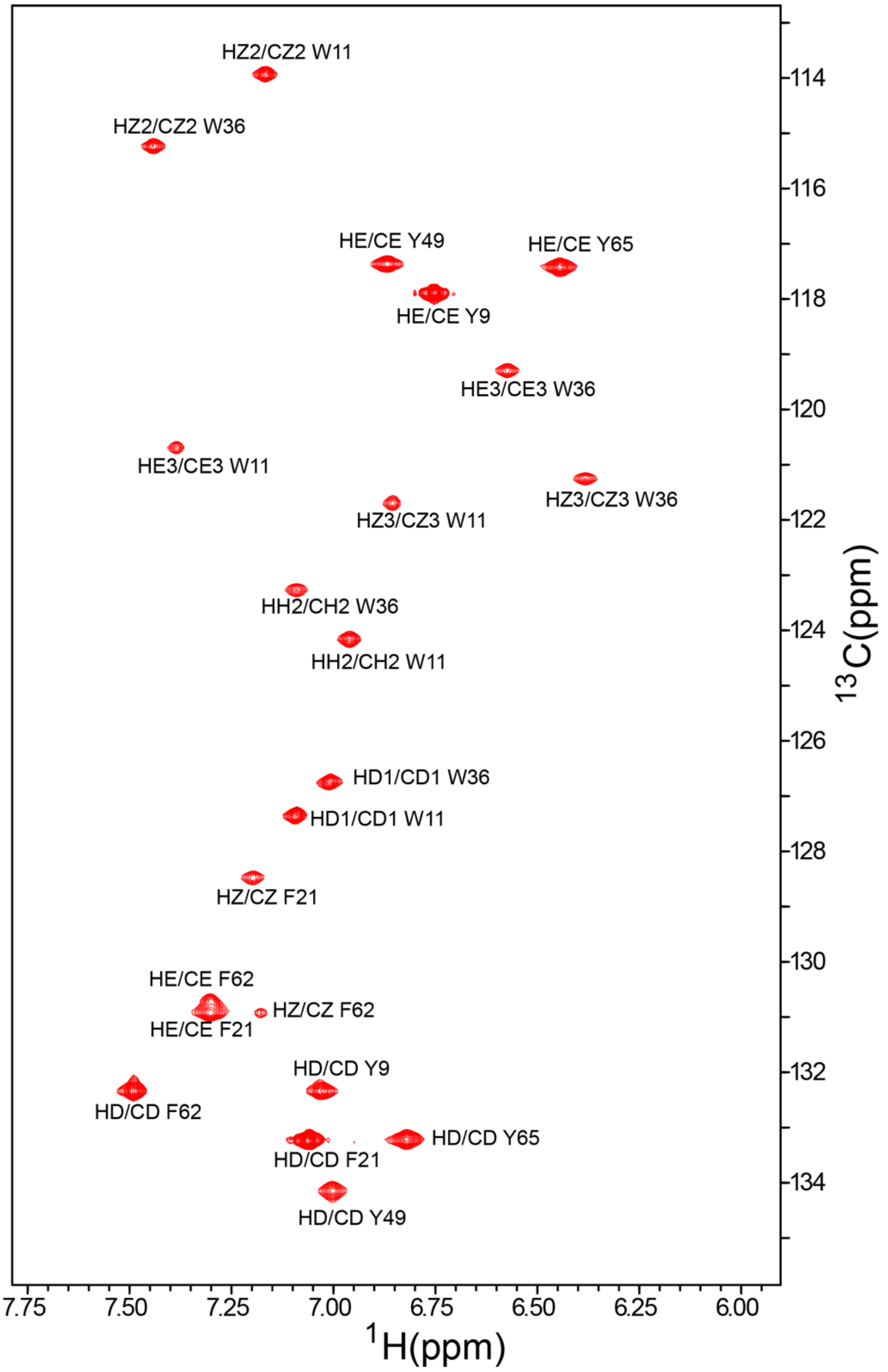

**Figure S10: The Fid-Net-2 processed aromatic $^{13}$C-$^{1}$H correlation map of the A39F FF domain (700 MHz; 274 K).** Assignments were obtained using the $^{1}$H-$^{13}$C-$^{1}$H NOESY-HSQC and $^{13}$C-$^{13}$C-$^{1}$H HSQC-NOESY-HSQC spectra.

### Supporting References

1. G. Karunanithy, D. F. Hansen, FID-Net: A versatile deep neural network architecture for NMR spectral reconstruction and virtual decoupling. *J. Biomol. NMR*. **75**, 179–191 (2021).

2. H. Geen, R. Freeman, Band-selective radiofrequency pulses. *J. Magn. Reson.* **93**, 93–141 (1991).

3. D. Marion, M. Ikura, R. Tschudin, A. Bax, Rapid recording of 2D NMR spectra without phase cycling. Application to the study of hydrogen exchange in proteins. *J. Magn. Reson.* **85**, 393–399 (1989).
